# UnitRefine: A Community Toolbox for Automated Spike Sorting Curation

**DOI:** 10.1101/2025.03.30.645770

**Authors:** Anoushka Jain, Robyn Greene, Chris Halcrow, Jake A. Swann, Alexander Kleinjohann, Federico Spurio, Severin Graff, Alejandro Pan-Vazquez, Björn Kampa, Juergen Gall, Sonja Grün, Olivier Winter, Alessio Buccino, Matthias H. Hennig, Simon Musall

**Affiliations:** Systems Neurophysiology, Department of Neurobiology, RWTH Aachen University, Aachen, Germany; Institute of Biological Information Processing (IBI-3), Forschungszentrum Jülich; Institute for Adaptive and Neural Computation, School of Informatics, The University of Edinburgh, UK; Centre for Brain Discovery Sciences, Hugh Robson Building, University of Edinburgh, UK; Department of Neuroscience, Physiology and Pharmacology, University College London, London, UK; Institute of Advanced Simulations (IAS-6), Forschungszentrum Jülich; Institute of Computer Science, University of Bonn; Princeton Neuroscience Institute, Princeton University, Princeton, NJ, USA; JARA BRAIN Institute of Neuroscience and Medicine (INM-10), Forschungszentrum Jülich, Jülich, Germany; Lamarr Institute for Machine Learning and Artificial Intelligence, Technical University Dortmund; Theoretical Systems Neurobiology, RWTH Aachen University; Champalimaud Foundation, Lisboa, Portugal; Allen Institute for Neural Dynamics, Seattle, WA, USA; Institute of Experimental Epileptology and Cognition Research, University of Bonn, Bonn, Germany

## Abstract

High-density electrophysiology simultaneously captures the activity from hundreds of neurons, but isolating single-unit activity still relies on slow and subjective manual curation. As datasets keep increasing, this poses a major bottleneck in the field. We therefore developed UnitRefine, a classification toolbox that automates curation by training various machine-learning models directly on human expert annotations. Fully integrated in the SpikeInterface ecosystem, UnitRefine combines established and novel quality metrics, cascading classification and comprehensive hyperparameter search to provide optimized models for different applications. UnitRefine achieves human-level performance across diverse datasets, spanning species, probe types, and laboratories, including recordings from mice, rats, mole rats, primates, and human patients. Applied to a large brain-wide dataset, UnitRefine doubled single unit yield and improved behavioral decoding performance. A streamlined graphical interface allows models to be fine-tuned to new datasets and shared via the Hugging Face Hub, enabling broad adoption and community-driven improvement of automated curation workflows.

## Introduction

Extracellular electrophysiology is a fundamental technique in neuroscience, enabling researchers to explore how individual neurons and neural populations encode and transmit information. Each recording electrode can capture action potentials from multiple individual neurons, necessitating the isolation and clustering of spiking activity through spike sorting. The goal of spike sorting is to first identify signals in the recording that represent putative action potentials, and then cluster them into groups, representing the activity of individual neurons (single-units)^1^.

Traditional spike sorting techniques rely on manual sorting and curation, yielding reliable results for individual electrodes or low-density probes such as tetrodes^2^. However, with the increasing number of recording sites on modern devices^3^, automated spike sorting solutions have been developed to improve the scalability and reliability of spike sorting while reducing the need for manual curation. These developments are critical for high-density multi-electrode arrays (HDMEAs) with hundreds or thousands of electrodes, where manual approaches are infeasible^4^. Large-scale datasets from recordings of brain-wide neural activity during different behaviors are now also openly available, establishing HDMEA recordings as a quasi-standard for many efforts to study neural circuit function^5,6^.

The significant volume of data generated by HDMEAs is a substantial challenge for scalable data processing and analysis^7,8^. Although automated spike sorters continuously improve, current algorithms still cannot reliably isolate single-unit activity^9,10^. Different experimental settings also introduce various sources of variability that can strongly affect spike sorting performance. For example, combining electrophysiology with optogenetic manipulation or functional imaging can introduce light-induced artifacts that can occlude spiking activity^11^. Moreover, movement artifacts are common in acute recordings and during behavior. This can limit the quality of single-unit isolation, and, in turn, affect the subsequent interpretation of neural recordings.

A potential solution for automated spike sorting curation lies in using metrics that quantify the quality of single-unit identification^12,13,14,15,16^. These metrics can be derived from features of action potential waveforms, measures of cluster separation, or from biophysical properties of action potential generation, such as refractory period (RP) violations. Manually setting thresholds on relevant metrics can then be used to automatically exclude noise or poorly isolated units. However, in many cases the distribution of the computed metric values varies depending on experimental conditions such as the recording system, recorded brain areas or species^17^. While this complicates identification of appropriate thresholds, it has been shown that an appropriate selection of metrics generalizes well across brain areas and recordings in mice from different labs^18^. Recent work has attempted to automate the curation progress^19,20^. However, existing approaches still require manual fine-tuning and expert knowledge to adjust for variability across recordings, and are primarily focused on Neuropixels recordings.

To overcome these limitations and enable fully automated spike sorting curation, we developed UnitRefine, a flexible open-source framework for the classification of spike clusters that is fully integrated into the Python-based open source toolbox SpikeInterface^10^. Users can train their own curation models on labeled data, or use existing, pretrained models. Using manually labeled data, we trained and optimized classifiers based on a large set of quality metrics computed from the clustered units. This approach yielded high classification accuracy in unseen recordings, comparable to the accuracy of expert human curators. Moreover, UnitRefine generalizes very well across different experimental settings, with high classification accuracy across various recording devices, including Neuropixels probes, Utah arrays, or wire bundles, and different species, including mice, rats, mole rats, monkeys, and humans. UnitRefine also outperformed current threshold-based approaches and improved the identification of task variables across brain regions, demonstrating its wide applicability and utility to accelerate neuroscientific discovery.

## Results

### Evaluation of manually curated spike-sorting clusters

To assess the quality of existing spike-sorting approaches, we analyzed a large set of acute mouse recordings obtained with Neuropixels 1.0 probes^21^, targeting different cortical and subcortical structures (Figure 1A, left). The overall recording quality varied substantially across sessions, with differences in baseline noise and spike visibility (Figure 1A, right). In some experiments, optogenetic stimulation or functional imaging introduced light-induced artifacts that degraded signal quality.

**Figure 1:**
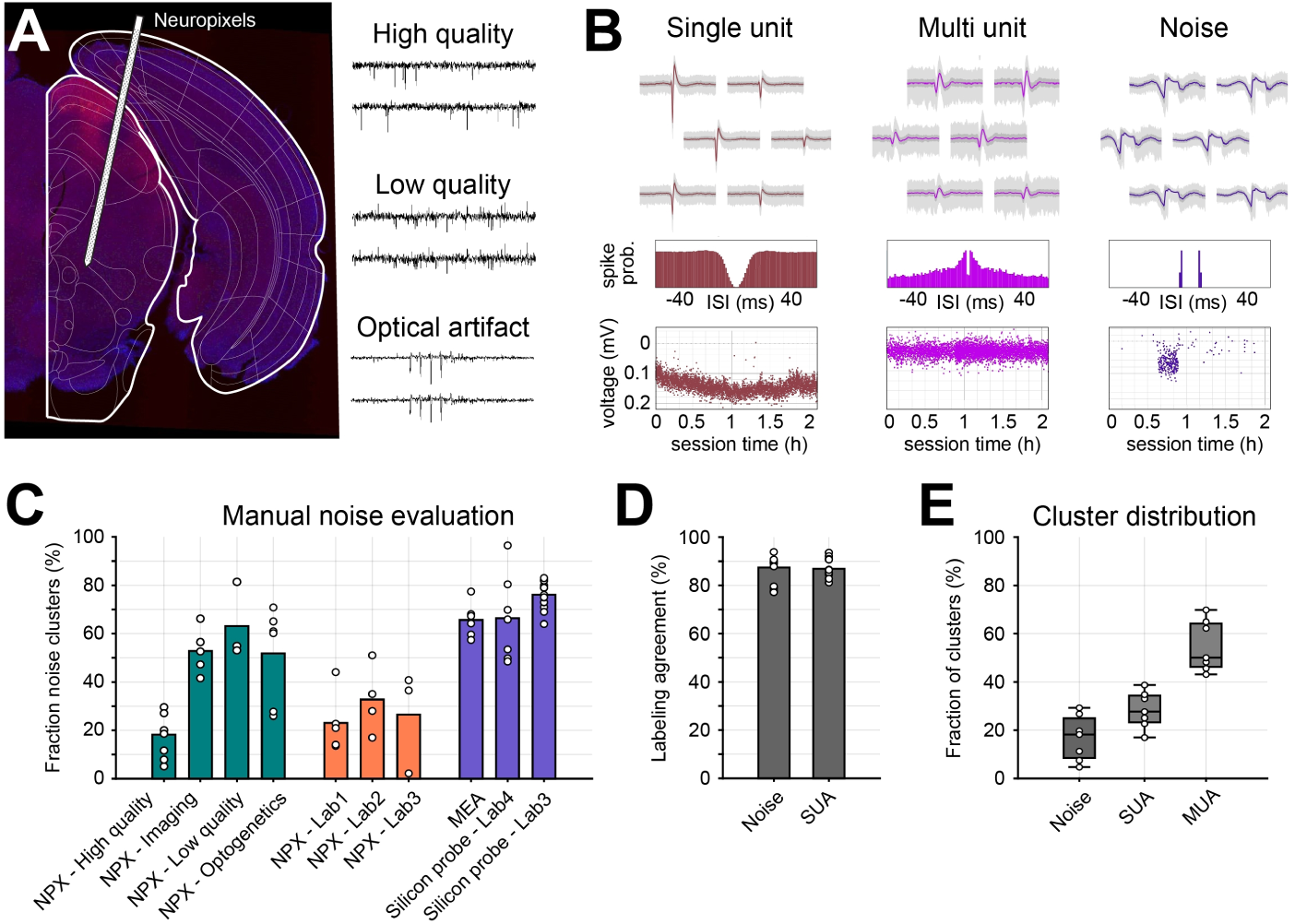
Multi-curator evaluation of different electrophysiological datasets. **A)** Left: Example brain slice with placement of a Neuropixels (NPX) probe across multiple brain regions. Right: Example traces for high- and low-quality recordings and optical artifacts. **B)** Example clusters for single-unit (SUA), multi-unit (MUA), and noise clusters. Top row shows the spike waveforms of each cluster across 6 recording channels. Middle row shows cross-correlogram of inter-spike intervals (ISIs) and bottom row shows the amplitude of all detected spikes across the entire recording. **C)** Manual evaluation of spike sorted clusters shows the fraction of noise clusters (%) across different recording types: 1) NPX of high or low quality and with additional functional imaging or optogenetic stimulation (green). 2) NPX recordings from other laboratories (orange), and 3) silicon probe and MEA recordings (purple). **D)** Labeling agreement across human curators for either separating noise from neural and MUA from SUA clusters (n = 11 recordings). **E)** Distribution of cluster labels, showing the fraction of noise, MUA, and SUA labels across all high-quality recordings (n = 7 recordings). Error bars indicate median ± 95% confidence intervals; individual recordings are shown as circles.

The base dataset was comprised of 11 Neuropixels 1.0 recordings containing 5,121 clusters from the primary visual cortex (V1), superior colliculus (SC), and anterior lateral motor cortex (ALM). A subset of seven high-quality recordings (2,945 clusters), consisted of homogeneous low-noise V1 and SC recordings and was used to develop and evaluate the classifier model (see below). Including four additional optogenetic recordings in the base dataset increased experimental diversity and exposed curators to a broader range of cluster types, further improving the robustness of the model between datasets (Supplementary Figure S1).

For spike sorting, we first isolated active channels on the probe, removed common-mode noise across channels using time-shifted median subtraction, applied a high-pass filter, and extracted putative single neurons with Kilosort 2.5^22^. The resulting clusters were manually labeled by five experienced curators using Phy^1^, following a standardized protocol that classified clusters as single-unit activity (SUA), multi-unit activity (MUA), or noise (Figure 1B).

SUA clusters were identified using three criteria: 1) Consistent spike waveforms across the cluster, with minimal variability and negligible interference from other waveform shapes. 2) A smooth correlogram of inter-spike-intervals (ISIs) with a clear dip in the center. 3) Stable spike amplitudes throughout the recording session with limited dispersion (Figure 1B, bottom row). MUA clusters contained neural but overlapping waveforms with lower amplitude and less variation across recording sites, and a larger number of RP violations due to contributions from multiple neurons. Noise clusters were characterized by irregular waveforms, ISI correlograms lacking a visible RP, and inconsistent spike amplitude distributions throughout the recordings.

Manual curation revealed a high fraction of noise clusters, particularly in low-quality or combined optophysiological recordings (51.7-52.8% mean noise occurrence). However, even in high-quality recordings, approximately 20% of all clusters were due to non-neural noise. This was also found in Neuropixels data from other research groups (23.1-32.7% mean noise occurrence; Figure 1C, orange), demonstrating that this issue is common across labs. Moreover, the proportion of noise clusters was even larger for other recording methods, such as multi-channel silicone probes or in vitro multi-electrode array (MEA) recordings (65.6-76.0% mean noise occurrence; Figure 1C, purple).

To assess the consistency of curator labels, we computed the cross-curator labeling agreement from two to five expert curators that independently annotated each recording in the base dataset. Across all recordings, we found high inter-curator reliability for noise and SUA (Agreement_Noise_ = 87.9% [77.9 − 89.9], Agreement_SUA_ = 86.4% [84.5 − 91.0]; median [95% CIs]) (Figure 1D). Although manual curation can show substantial variability across experts^23,10^, this shows that our human curators consistently labeled clusters with high accuracy, providing an important basis to develop an automated curation approach. The overall distribution of labels across high-quality recordings is shown in Figure 1E. All annotated datasets are also publicly available^2^.

### Novel cluster metrics for automated spike-sorting curation

After manually labeling the spike-sorted clusters, we evaluated how well different quantitative metrics could predict human labels. We therefore used SpikeInterface to compute a wide range of cluster metrics^3^, and split them into quality and template metrics. Quality metrics assess the spike sorting reliability and contain various features of spike trains, spike amplitude, and overall cluster separation^15,24^. Template metrics capture neural waveform properties that distinguish neural activity from artifacts, such as the peak-to-valley duration or full-width half maximum^4^. A detailed description of all metrics can be found in the appendix.

We also developed three new cluster metrics to address specific noise and SUA features and implement them in SpikeInterface. Synchrony metrics quantify the simultaneous occurrence of spikes across multiple clusters, leveraging spike synchronicity to identify noise due to electrical interference or channel crosstalk^25^. To account for this feature, we established two separate measures, quantifying the number of simultaneous spikes occurring within at least two or four different clusters.

Amplitude variation metrics quantify the stability of spike amplitudes over time, which should be high for well-isolated neurons. Amplitude variation is based on the ratio between the standard deviation and the mean of spike amplitude, computed at various recording times, from which the amplitude variation median and range are computed.

Lastly, firing range quantifies the firing rate dispersion over time. We first computed the firing rate distribution within 10-s long recording periods and defined the firing range as the the difference between the 5th and 95th percentile of the local firing rate distribution. Very high variations in firing range are characteristic of noise clusters which tend to exhibit large dispersion at specific recording times (Figure 1B, bottom right panel).

### Discriminative power of individual cluster metrics

We next evaluated the capacity of individual metrics to predict cluster quality and whether they provided complementary information. Pairwise correlations showed that metrics were only weakly correlated with few expected exceptions, such as a high correlation between the geometric median spike amplitude and signal-to-noise ratio (Figure 2A). Principal component analysis (PCA) also showed that the explained variance gradually increased, with 22 components required to account for 90% variance (Figure 2B). Different metrics therefore, carry complementary information, motivating their combined use for cluster curation.

**Figure 2:**
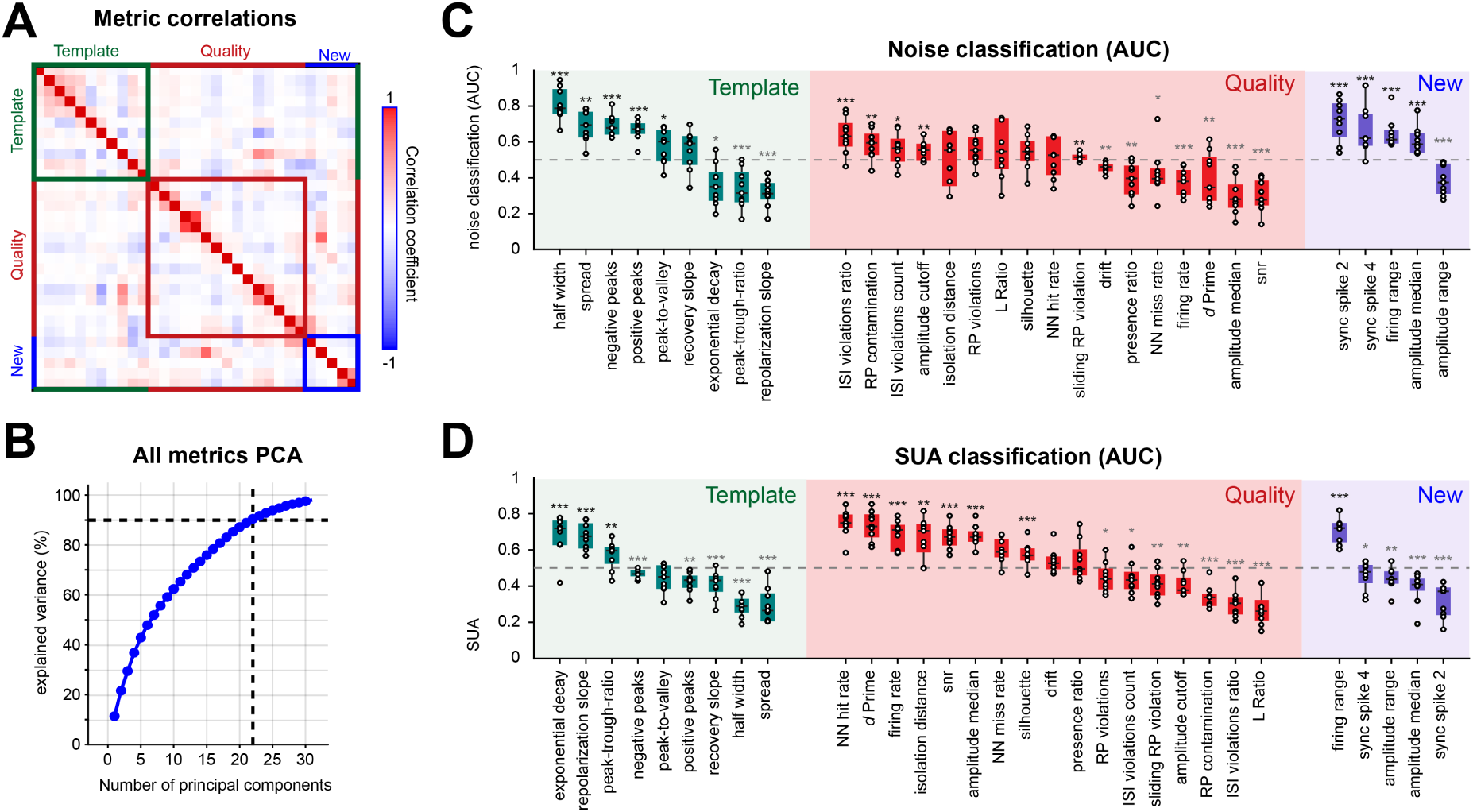
Discriminative power of neural quality metrics. **A)** Correlation matrix showing relationships between all metrics. Red and blue indicate positive and negative correlations, respectively. **B)** Cumulative explained variance from principal component analysis, showing that 22 components are required to explain 90% of variance (dashed lines). **C)** AUC values for distinguishing noise from non-noise clusters, using established template metrics (green), quality metrics (red), and new quality metrics (purple). AUCs near 1 or 0 indicate strong predictive power; values near 0.5 indicate no relationship. Significance was assessed using bidirectional Wilcoxon signed-rank tests (\*\*\**p <* 0.001, \*\**p <* 0.01, \**p <* 0.05). **D)** Same as in C), but for SUA classification. Box plots show the median (central line), interquartile range (box), and 95% confidence intervals (whiskers). Circles indicate individual recordings (n = 11). Metrics are ordered by descending median AUC.

To assess the information content of individual metrics over human labels we used an area under receiver operating characteristic (AUC) analysis^26^. Almost all metrics yielded significant predictions for noise classification (AUC significantly different from 0.5, Wilcoxon signed-rank test, *p <* 0.05) with the half-width of the cluster waveform and the synchrony metrics being particularly informative (Figure 2C). Most metrics could also significantly discriminate between MUA and SUA clusters (Figure 2D) with inter-spike-interval violations, L ratio, firing range and synchronicity being the most informative.

Together, these results show that metrics are complementary and differentiate between noise, MUA, and SUA, suggesting that combining a larger group of complementary metrics into a decoder framework could yield high automated curation accuracy.

### Development and evaluation of an automated curation approach

To develop a fully automated curation framework, we evaluated two classification strategies. In the cascaded approach, clusters were first divided into noise and neural groups, followed by cluster separation into MUA and SUA. Alternatively, SUA clusters were classified directly against all non-SUA clusters. Classifier hyperparameters were optimized using a Bayesian search procedure, and performance was evaluated with leave-one-session-out 5-fold cross-validation. All reported accuracies are therefore based on classification of completely unseen recordings.

Using all computed metrics from the high-quality base dataset, we found a consistently strong performance for the separation into noise and neural clusters, and subsequently into SUA and MUA (Figure 3A,B,D,E; Table 1, 2). Among the tested models, ensemble methods performed best, with RF classifiers yielding the highest balanced accuracy (Accuracy_Noise_ = 83% [73.8 − 90.9], Accuracy_SUA_ = 79.3% [70.7 − 85.6]; median [95% CIs]). To assess how performance scales with dataset size, we examined learning curves across classifiers (Figure 3C and 3F). Balanced accuracy increased rapidly with training data and gradually plateaued after approximately 500 clusters. Thus, curated datasets from one or two Neuropixels recordings could already be sufficient for robust decoder training, with further data providing more gradual improvements.

**Figure 3:**
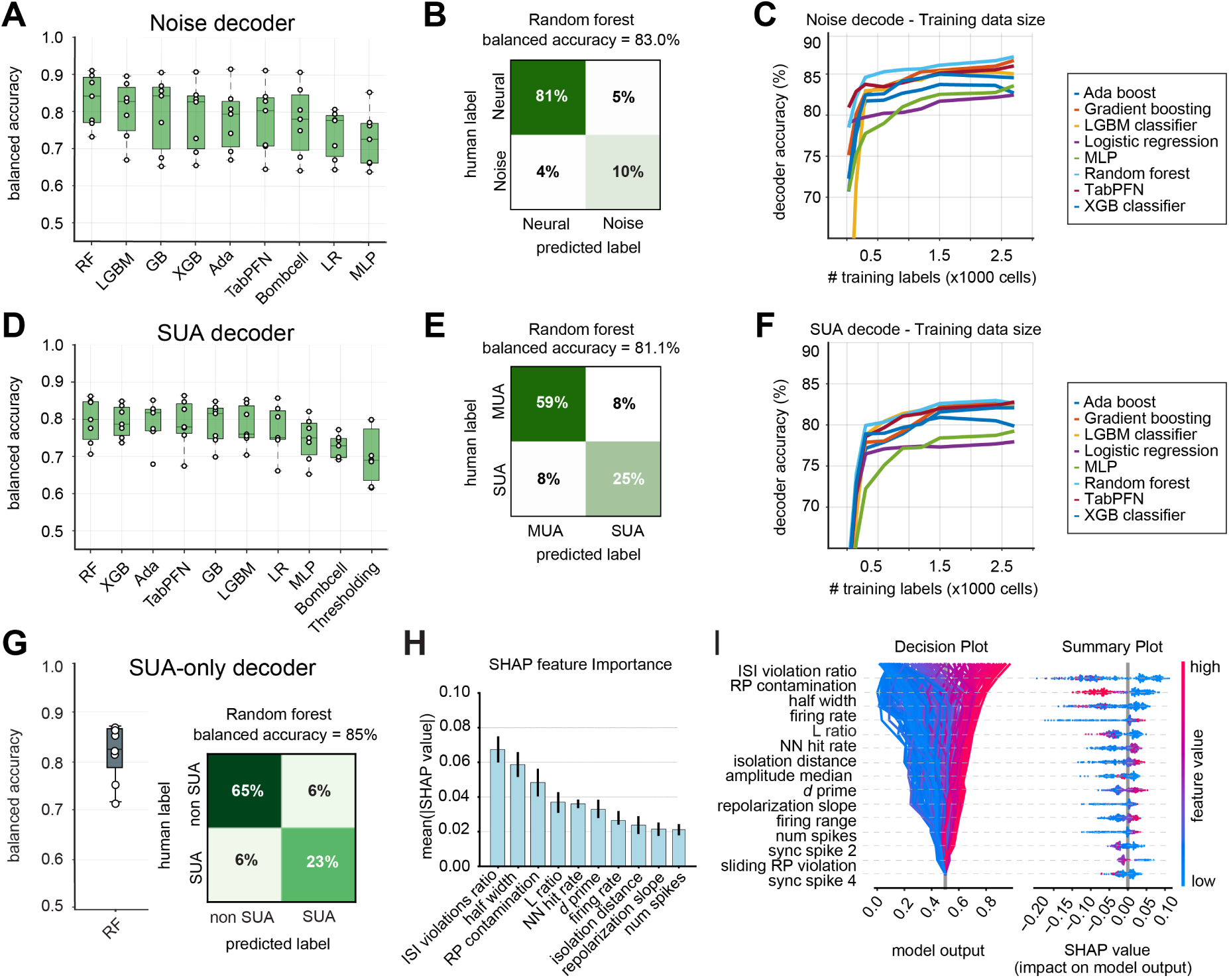
Comparison of machine learning classifiers for noise and SUA separation. **A)** Comparison of balanced accuracy across different classifiers for noise decoding. Ensemble methods consistently outperformed linear models, neural networks, and the transformer-based TabPFN. **B)** Confusion matrix for the RF model, showing an overall balanced accuracy of 83%, with equally low false positive and false negative rates. **C)** Learning curves demonstrate how classifier performance scales with increasing training data for noise classification. RF reached high accuracy with relatively few labeled clusters, stabilizing around 500 clusters. **D–F)** Same as in A–C) for SUA classification. Ensemble methods again achieved the highest performance, with RF showing rapid gains and stable accuracy across datasets. **G)** Left: Balanced accuracy of the Random Forest classifier for distinguishing SUA from all other clusters. Right: Confusion matrix, showing low misclassification of cluster labels. **H)** SHAP-derived feature importance across leave-one-out validations for SUA classification for the 10 most important metrics. Unit quality metrics (e.g., ISI violation ratio, L-ratio, RP contamination) consistently contributed the most to model predictions. **I)** SHAP-based quantification of RF predictions for SUA classification, visualized for 200 randomly sampled clusters from an example recording for the 10 most important metrics. The decision plot (left) depicts the influence of features on individual model predictions, while the summary plot (right) shows the overall importance and directionality of each feature across the dataset.

We also tested curation with Bombcell (v1.7.2), a popular threshold-based method for cluster classification optimized for Neuropixels data^20^. After manual optimization, Bombcell achieved a balanced accuracy of 77.0 [69.2 − 86.1]% for noise and 72.6 [69.2 − 76.9]% for SUA detection.

Notably, SUA classification showed very high precision (few false positives) but lower recall (more false negatives; Supplementary Figure S2). Similar results were observed when using a combination of recommended metrics thresholds from the Allen Institute for Neural Dynamics^5^ (Figure 3D). Together, these results highlight an inherent trade-off: conservative thresholding reliably identifies SUA clusters but excludes potentially useful clusters, whereas machine-learning classifiers emulate human curator behavior to achieve higher sensitivity.

We also tested a classifier to directly isolate SUA from all other clusters, as commonly done in other datasets^27^. The RF model again achieved high performance, with 85% balanced accuracy across seven recordings containing 836 SUA clusters (28.4%) (Figure 3G). Overall, model performance approached the upper bound of inter-curator agreement for SUA labeling (median 86.4%, 95% CI: 84.5–91.0%; Figure 1D), demonstrating its reliability for automated curation of unseen recordings.

To identify which features determine SUA classification, we applied cross-validated Shapley additive explanations (SHAP^28^) to the RF model, the most accurate classifier in our evaluation (Figure 3H, Supplementary Figure S3). For each cluster, SHAP assigns one value per metric, indicating how much that metric shifts the prediction toward SUA (positive values) or toward non-SUA (negative values). ISI violations ratio was the most discriminative metric, followed by spike half width, and RP contamination with highly consistent feature rankings across training subsets. The decision plot for an example recording illustrates the contribution of each feature, with individual clusters shown as trajectories toward their classification outcome (Figure 3I). SHAP values were typically bimodal, with large metric values consistently shifting predictions toward non-SUA classifications. These results show that even with limited training data, the classifier leverages biologically meaningful metrics and performs curation in a manner comparable to human experts.

### UnitRefine generalizes across labs and brain regions

To assess cross-dataset generalization, we tested the pre-trained classifier from our lab on independent recordings from multiple laboratories, targeting distinct cortical and subcortical brain regions.

We first evaluated our pre-trained model on forebrain recordings from the IBL dataset (8 sessions; see Table 4)^29^, achieving robust balanced accuracy of 79% (Figure 4A). SHAP-based feature attribution also revealed largely consistent metrics across datasets, except median amplitude having a surprisingly strong impact on model decisions (Figure 4B). Single-cluster SHAP values for median amplitude were also predominantly negative, thus shifting the model toward non-SUA classifications (Figure 4C). This suggests that cross-dataset differences in spike amplitude may have reduced curation performance. Indeed, while the pre-trained model clearly outperformed threshold-based curation using the IBL standard criteria^30^, retraining the model directly on IBL data further improved performance, reaching a very high balanced accuracy of 98% (Figure 4D, E). Although the pre-trained model readily generalized to new recordings, retraining therefore further aligned it to dataset-specific recording conditions. After retraining, ISI violation ratio, firing rate, and RP contamination emerged as the most informative features (Figure 4F). Notably, the mean absolute SHAP value for ISI violation ratio increased from 0.060 in the pre-trained model to 0.150 after IBL retraining (Figure 4B, F), reflecting its heightened contribution to classification accuracy. This shows that although key physiological features remain consistent across datasets, their relative weights also adapt to lab-specific curation practices.

**Figure 4:**
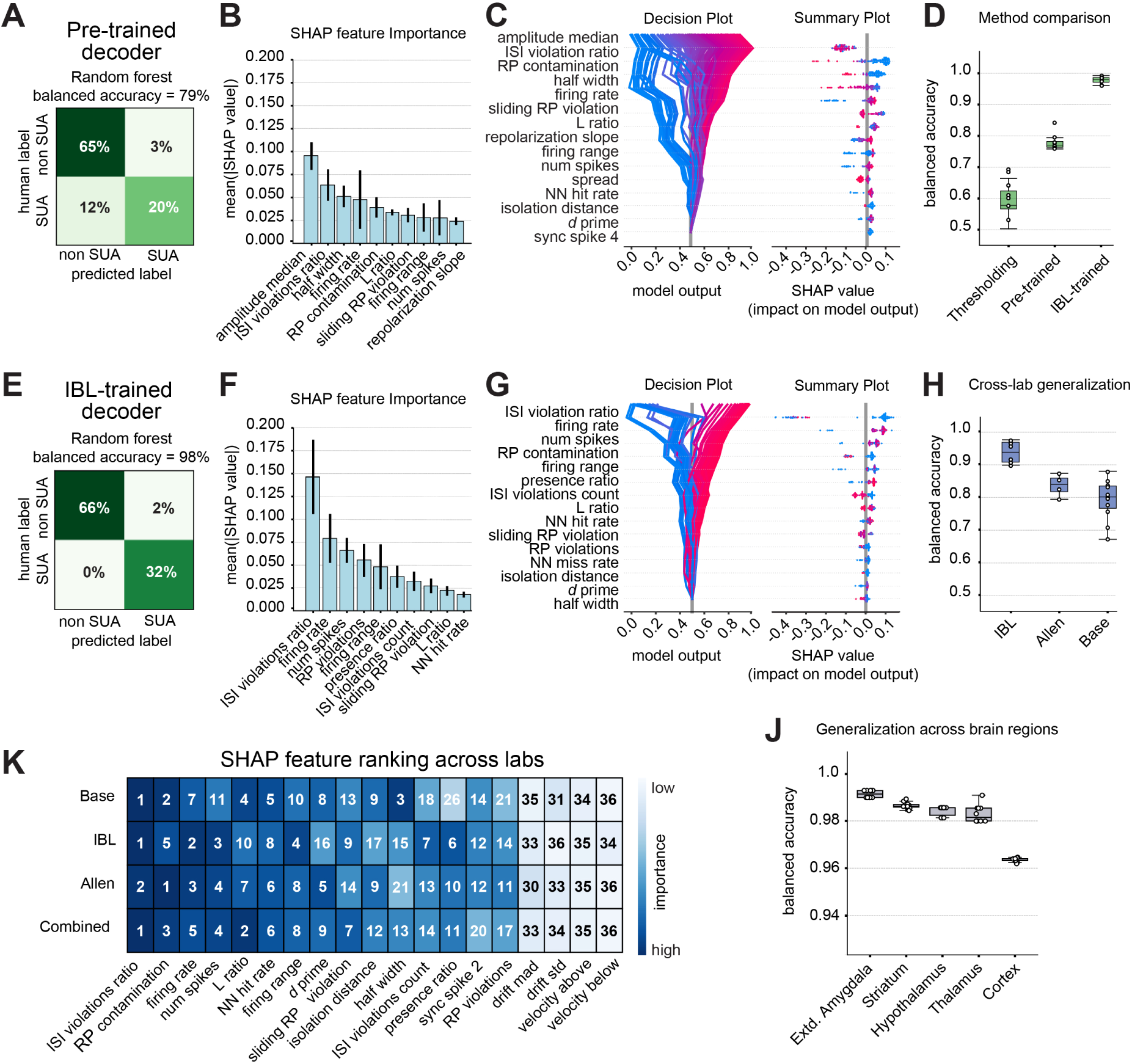
Cross-laboratory and cross-region generalization of automated curation. **A)** Confusion matrix showing 79% balanced accuracy for a pre-trained RF model, tested on IBL recordings. **B)** SHAP-derived feature importance across leave-one-session-out validation for the pre-trained model, highlighting amplitude median, ISI violation ratio, and RP contamination as important metrics. **C)** SHAP-based quantification of pre-trained model predictions, visualized for 200 randomly sampled clusters. The decision plot (left) depicts the influence of features on individual model predictions, while the summary plot (right) shows the overall importance and directionality of each feature across the dataset. **D)** Leave-one-session-out validation, comparing balanced accuracy with thresholding, pre-trained, and IBL-trained models. **E-G)** Same as in A-C) for an IBL-trained model. **H)** Balanced accuracy of the unified cross-lab model, using leave-one-session-out validation across the base, IBL, and Allen datasets. **I)** Ranking of SHAP feature importance across labs, highlighting both universally predictive quality metrics, such as ISI violations ratio, and lab-specific variations in waveform metrics, such as spike half-width. **J)** Balanced accuracy across five major brain regions (cortex, extended amygdala, hypothalamus, striatum, thalamus) for IBL data, showing robust generalization (*>*95% accuracy) across 10 random seeds.

We next tested the model on Neuropixels 2.0 recordings from the Allen Institute for Neural Dynamics (4 sessions; see Table 4), which provide a denser electrode layout than Neuropixels 1.0 probes, and observed consistent performance (Supplementary Fig. S4).

Here, the pre-trained model achieved high balanced accuracy of 86%, with a modest improvement to 91% when retraining on Allen sessions, demonstrating robust cross-dataset generalization. To determine whether a cross-laboratory training strategy further improves model generalization, we pooled all available recordings (*n* = 23) and trained a unified RF model. Leave-one-session-out validation revealed consistently high balanced accuracy across all datasets and labs (Figure 4H). This shows that pooling data from different labs and recording modalities can improve model robustness and mitigate lab-specific biases, thus reducing the need for retraining on individual datasets. All classifiers and the foundational datasets are also available on Hugging Face^5^. Comparing SHAP feature rankings of the unified model across datasets revealed that quality metrics, such as ISI violation ratio, RP contamination, and firing rate, were consistently among the most informative features (Figure 4K). In contrast, waveform metrics, such as spike half-width, were more variable, suggesting that spike waveform properties are more dependent on recording-specific parameters, such as probe type or recording depth. Lastly, we also evaluated curation performance across different brain regions in the IBL recordings, including the frontal cortex, extended amygdala, hypothalamus, striatum, and thalamus. Region-wise benchmarking showed high balanced accuracies (96–99%, Figure 4J), demonstrating robust generalization across distinct brain areas.

### Evaluating metric subset importance for model generalization

Having established that UnitRefine achieves robust performance across multiple labs and brain regions, we asked if the model maintains high accuracy with a reduced subset of core metrics. Given the diversity of standards across electrophysiology systems, this is particularly important when using a generalized model on recordings where some specialized metrics might not be available.

We therefore evaluated model performance across all datasets (*n* = 23 recordings, 10,833 clusters) using four different groups of metrics that balanced physiological relevance, availability, and computational efficiency. Group 1 consisted of all metrics; Group 2 contained only quality metrics; Group 3 contained spike time-based metrics, such as ISI violations and spike synchrony, which are rapidly computed from spike timestamps alone; and Group 4 contained only template metrics related to waveform morphology (Table 3).

As expected, classification accuracy was highest when using all metrics (Figure 5A, Table 5). Notably, spike time-based metrics yielded nearly equivalent balanced accuracy at minimal computational cost, making them particularly valuable for resource-limited settings. In contrast, quality metrics produced slightly lower accuracy, and template metrics alone were least effective. Thus, spike time metrics provide an efficient foundation for a generalized curation model that can be translated to any recording system. We therefore used spike-time based models to test UnitRefine across species.

**Figure 5:**
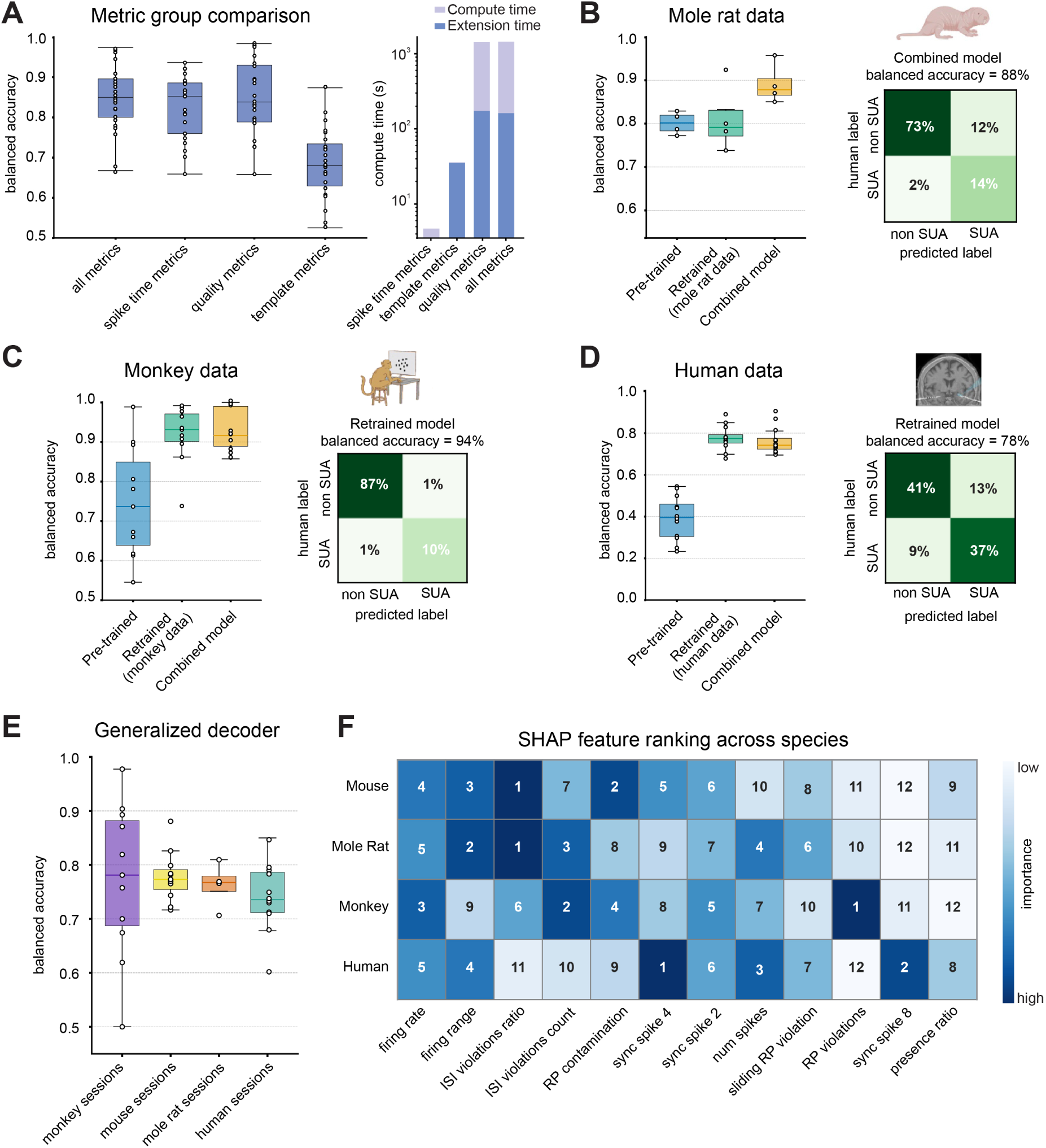
Cross-species generalization of UnitRefine. **A)** Left: Leave-one-out balanced accuracy across classifiers, trained on different metric groups. Right: Compute time for an example 25-minute Neuropixels recording, split into feature extraction (blue) and cluster metric computation time (purple). **B–D)** Species-specific leave-one-out balanced accuracy of different classifiers for mole rat (**B**), monkey (**C**), and human (**D**) datasets. Boxplots show balanced accuracy across recordings; confusion matrices illustrate correspondence between human and predicted labels. **E)** Leave-one-out balanced accuracy of a generalized classifier, applied to datasets from different species. Despite some variability, the classifier achieved high median accuracy across all species. **F)** SHAP-based ranking of feature importance across species. Universally informative metrics include ISI violation ratio, RP contamination, and firing rate, while amplitude- and waveform-based features vary across species.

### UnitRefine generalizes across recording systems and species

We first examined cross-species generalization in mole rat recordings (4 sessions; see Table 4). The mouse-trained model already achieved high balanced accuracy, suggesting effective transfer of spike-based metrics across species. Retraining a mole rat–specific model yielded similar performance, while a mixed-species model, combining mouse and mole rat data, achieved the highest balanced accuracy (Figure 5B).

Augmenting the existing rodent model with mole rat data, therefore, strengthened decoder reliability, providing a valuable alternative to retraining from scratch. Moving beyond rodents, we applied UnitRefine to nonhuman primate recordings (11 sessions). The mouse-trained model achieved moderate curation accuracy, while retraining on monkey data substantially improved performance. Notably, a mixed mouse–primate model achieved comparable accuracy while expanding training diversity (Figure 5C), indicating robust transferability across species and recording systems.

Finally, we applied UnitRefine to human intracranial recordings (12 sessions). The mouse-trained model performed poorly, whereas retraining on human data strongly improved accuracy, with equally high performance of a mixed mouse-human model (Figure5D).

Across all species, mixed models consistently performed comparably to species-specific decoders, highlighting the benefit of combining annotated training datasets from different species. A unified cross-species model also achieved high leave-one-session-out performance across all species, demonstrating that it captured core electrophysiological patterns that generalize beyond individual datasets and species-specific biases (Figure 5E). These results underscore that UnitRefine can be flexibly extended to unseen species, recording platforms, and experimental conditions with the unified cross-species model providing a versatile starting point for diverse electrophysiological experiments.

SHAP-based feature importance ranking showed that firing rate was the most consistent metric across species, while ISI violation ratio was among the most informative metric in all animal models. In contrast, human data show a heavier reliance on synchrony-based metrics, potentially due to differences in electrode geometry and noise structure. Together, these findings demonstrate that UnitRefine maintains robust generalization across species, probe types, and spike-sorting pipelines, supporting its use from basic to translational neuroscience applications.

### UnitRefine improves cluster yield and population analysis sensitivity

To demonstrate the practical impact of UnitRefine, we applied it to the large-scale brain-wide IBL dataset of 699 recordings from task-performing mice^27^. In the original study, SUA clusters were identified by thresholding amplitude, RP violations, and amplitude symmetry, reducing the 621,733 clusters to 75,708 (12.8%)^30^. In contrast, UnitRefine identified 175,027 (28.1%) putative SUA clusters, which substantially overlapped with threshold-selected clusters (87.9%) while greatly expanding the total amount of available clusters (Figure 6A). Next, we compared population-level decoding performance using UnitRefine- or threshold-selected clusters to predict animal choices and feedback outcomes across brain regions. To ensure that the differences were due to the principled selection of informative units rather than an increased sample size, we included a control condition containing all threshold-selected clusters, plus a random subset of additional clusters to match the number of UnitRefine clusters. While the control consistently underperformed in both groups, UnitRefine led to a modest but consistent increase in the accuracy of the feedback prediction (Figure 6B). More importantly, UnitRefine increased the fraction of subregions within each brain structure that were significantly predictive of animal choice or feedback, indicating increased sensitivity to detect task-related signals in individual brain areas (Figure 6C). The differences in task decoding between the two curation approaches were broadly distributed in general and strongest in the frontal cortical and midbrain areas for choice and the hindbrain areas for feedback (Figure 6D). Gains were particularly pronounced in smaller brain areas, such as the orbitofrontal cortex or midbrain reticular nucleus (Figure 6E), suggesting that an important advantage of UnitRefine curation lies in increasing the number of high-quality clusters when the unit numbers are otherwise too low for robust population analyzes.

**Figure 6:**
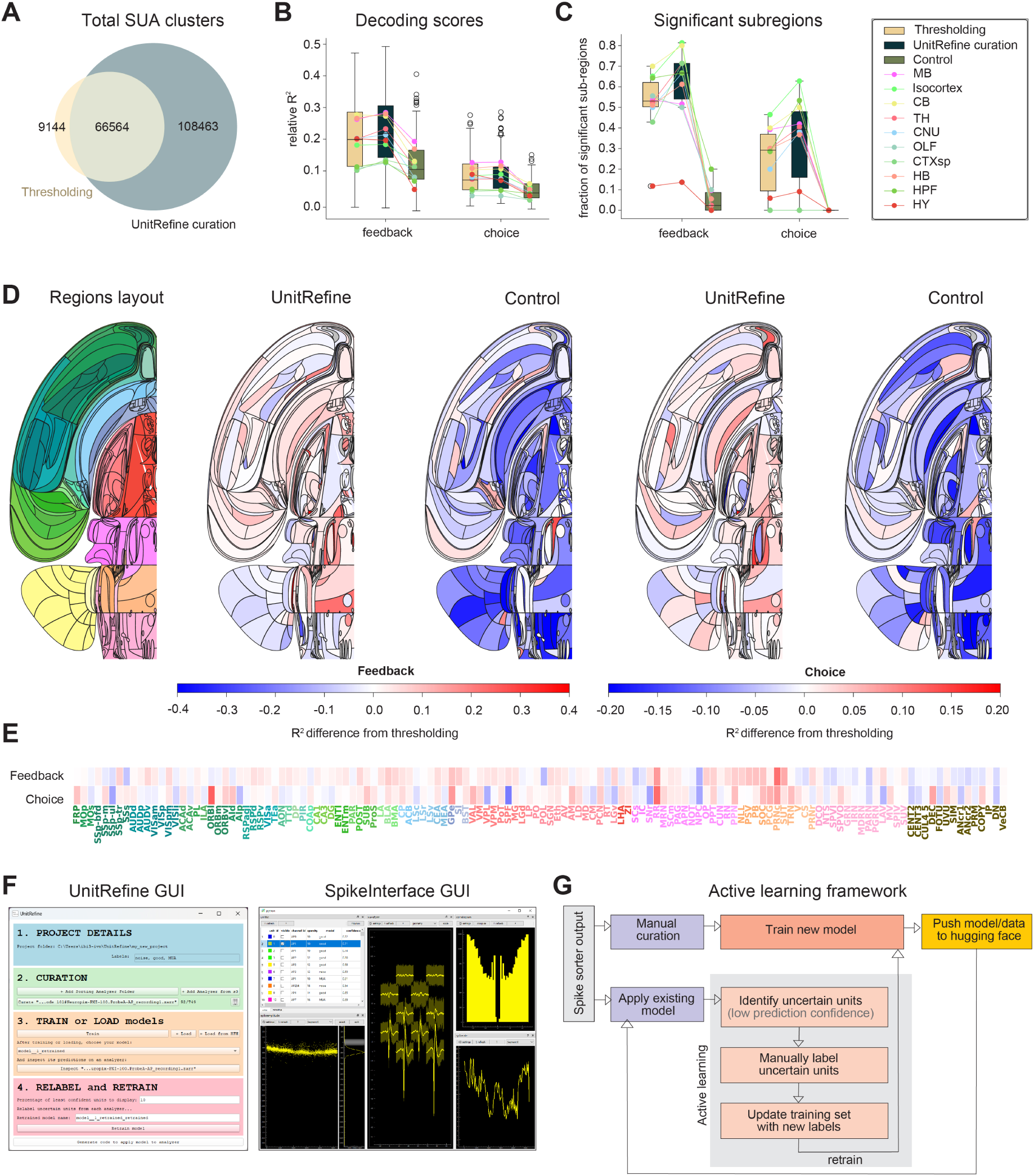
Impact of automatic curation strategy on a large-scale dataset. **A)** Venn diagram of SUA clusters in the IBL dataset, either based on metric thresholding or UnitRefine. **B)** Decoding performance for feedback and choice across brain regions, computed as the null-corrected median balanced accuracy of a logistic regression decoder. **C)** Fraction of subregions within each major brain structure that were significantly predictive of animal choice or feedback. **D)** Left: Swanson flat maps of the whole brain with regions color-coded by anatomical assignment. Additional information on brain regions is available online^6^. Right: Difference maps for feedback and choice decoding with UnitRefine or control curation versus thresholding curation. Areas with improved or decreased decoding performance are shown in red or blue, respectively. **E)** Same data as in D) but with individual area labels. Font colors indicate membership to brain regions. **F)** UnitRefine-GUI (left) illustrating the four main steps: data loading, cluster curation, model training or loading, and validation-based model retraining. Cluster curation is done through the SpikeInterface-GUI (right). **G)** Active-learning–based workflow for iterative model refinement. The existing model is applied to new data, uncertain and high-confidence units are presented to the user, and user feedback is used to retrain and improve model performance.

### Cluster curation and active learning

To promote broad adoption, we developed a graphical user interface (GUI) with detailed online doc-umentation^7^ to navigate the UnitRefine workflow (Figure 6F, left). Users can load data from local or cloud storage, train new models from curated labels, or apply existing UnitRefine models. In a subsequent validation step, predicted labels can be examined, corrected and also be leveraged to re-train customized models. Validation and curation is implemented through Spikeinterface-GUI, which is used to visualize key cluster features and improve the curation quality. An integrated active learning pipeline (Figure 6G) further enhances model customization by automatically highlighting clusters with low prediction confidence for user review. These additional labels are incorporated into the training set to iteratively improve model performance. Finally, trained models and datasets can be shared via the HuggingFace Hub, facilitating collaboration and reproducibility across labs.

## Discussion

Our results demonstrate that fully automated curation of spike sortings is feasible and achieves high accuracy comparable to that of human experts. Using a supervised learning framework based on established and novel quality metrics, UnitRefine accurately classifies neuronal units across diverse datasets and experimental conditions, spanning different brain areas, recording systems, and species. Designed as a flexible analysis pipeline, UnitRefine accommodates various preprocessing algorithms and spike-sorting methods while maintaining high accuracy in identifying SUA clusters that are essential for downstream analyses.

Automated curation addresses a critical bottleneck in current electrophysiology workflows: the extensive manual labor required to refine sorted clusters. By reducing or eliminating the need for expert intervention, UnitRefine can save hundreds of hours in human effort per lab, allowing researchers to focus on experimental interpretation rather than technical validation. In settings where manual curation remains desirable, UnitRefine labels can improve decision speed and consistency by highlighting uncertain clusters for review. Moreover, its active learning framework enables rapid retraining or creation of new models for custom applications. Combining predictions with traditional thresholding rules can also be used to enhance sensitivity. Such hybrid approaches may be particularly effective for large or noisy datasets, as well as for emerging applications like closed-loop BMI systems where real-time spike sorting is essential^31^.

A key strength of UnitRefine lies in its integration with SpikeInterface^10^, which standardizes metric computation, streamlines and simplifies the construction of analysis pipelines, and facilitates the reuse and sharing of trained models. In addition, we provide a standalone implementation for expert users to incorporate UnitRefine into custom workflows, allowing flexible integration of additional cluster metrics for specialized applications, such as automatic identification of neurons with specific tuning properties. The conceptual direction of this work aligns closely with ongoing efforts to modularize spike-sorting pipelines within the SpikeInterface framework. Traditional sorters are typically implemented as monolithic packages with multiple algorithmic stages, such as filtering, spike detection, feature extraction, and clustering^1,16,32^. However, performance often varies substantially across stages. A modular design allows individual components to be interchanged, promoting both flexibility and transparency in pipeline optimization. UnitRefine offers a principled way to benchmark each stage by evaluating how changes in preprocessing, peak detection or clustering methods affect the final curation quality. In the absence of real ground truth, predicted labels can serve as a proxy to identify the most reliable pipeline configurations for a given dataset. This could also be beneficial for future features, such as automatic cluster splitting or merging, that may be required for fully automated curation procedures.

With the growing availability of large standardized datasets, for example from the IBL^18^ and the Allen Institute for Brain Science^5^, including diverse datasets from individual laboratories will further enable researchers to evaluate how well spike-sorting methods generalize across different experimental conditions and assess their robustness to variations in recording setups and preprocessing strategies. While generalization across regions and species was overall robust, we found lower accuracy in some hindbrain and cerebellar regions (Figure 6D), likely because they were not contained in the training data. Especially for regions where spike waveforms are strongly shaped by anatomy, for example Purkinje cells or climbing fibers in the cerebellum^33^, region-specific models may therefore provide more accurate classification.

A major obstacle in developing novel SUA classification algorithms is also the absence of ground-truth datasets^9^. A notable exception is SpikeForest^34^, which combines real ground-truth datasets and simulated data to enable systematic benchmarking of spike sorters. Such standardized datasets are essential for reproducibility and meaningful comparison across different stages of the spike sorting process. To contribute to these efforts, all spiking data, cluster metrics, and human labels that we used in the study are therefore available in an online repository^35^.

The UnitRefine models trained on the base dataset used 37 distinct metrics per cluster. However, many of these metrics are computationally expensive because they depend on features that require scanning the entire raw recording, such as individual spike amplitudes and PCA analyses. Moreover, some metrics are specific to HDMEAs, limiting the transferability of models trained with them to other recording types. Fortunately, our model comparison showed that spike-time-based metrics alone can achieve very high curation accuracy across many recording types, making them ideally-suited for generalized models that can be employed to virtually any electrophysiological experiment (Figure 5). Both species-specific and generalized models are provided within the UnitRefine workflow to serve as a starting point for automated curation of new datasets. Spike-time metrics can also be computed directly from different spike-sorter outputs without requiring access to raw data, further broadening the applicability of the approach. In addition, only around 500 curated clusters were needed to train a robust decoder, suggesting that smaller laboratories can readily adapt UnitRefine by testing existing models, verifying predicted labels, and curating a limited number of clusters to further refine model accuracy. Iteratively improving the initial automated curation in this way quickly increases the number of available training clusters and enhances overall performance.

Retraining becomes particularly important when recording conditions differ substantially from the training data. For example, the base model performed poorly on human intracranial recordings, likely due to differences in the noise structure, electrode geometry or the used spike-sorter, but substantially improved after retraining. When applying UnitRefine to a new species or recording type that was not included in the training data, augmenting an existing generalized model with limited species-specific labels is therefore likely to yield the highest classification accuracy. Importantly, UnitRefine is also not limited to specific cluster labels and users may apply it to other settings, such as the automated classification of neuron subtypes^36^ or somatic versus axonal activity^20^. Threshold-based tools such as Bombcell can also provide complementary value by identifying dataset-specific criteria, and their labels can be directly used to train or refine UnitRefine models. We therefore believe that UnitRefine can be a highly valuable tool across a wide range of experimental settings that include electrophysiological recordings, promoting standardization while reducing manual labor.

Finally, developing UnitRefine as an open-source Python package ensures accessibility, transparency, and community growth. Its integration with SpikeInterface enables precise version control, reproducible benchmarking, and straightforward troubleshooting, which are essential for methodological rigor. By lowering technical barriers to automated spike curation, UnitRefine offers a reproducible, efficient, and user-friendly solution for modern electrophysiological analysis. We anticipate that its combination of performance, openness, and extensibility will promote widespread adoption, accelerating progress toward standardized, automated, and interpretable neural data processing across the neuroscience community.

## Methods

### Neural Recording Acquisition and Preprocessing

For Neuropixels 1.0 recordings in the base dataset, probes were stereotactically positioned in V1 and the midbrain through a thin silicone window using standard procedures^21^. In experiments involving optogenetic stimulation, light delivery was synchronized with the electrophysiological recording to monitor potential artifactual responses. Further surgical details for these recordings are also provided in Balla et al.^37^.

Raw electrophysiological data were preprocessed using temporal phase shifting, identification and removal of bad recording channels, bandpass filtering (300–6000 Hz), and common-mode referencing^8^. Transient artifacts were identified and removed with CatGT^9^. Spike detection and sorting were then performed with Kilosort 2.5^38^ using default parameters.

Neural recordings from different labs were acquired using multiple electrode and recording systems, including Neuropixels 1.0 and Neuropixels 2.0, and other silicon probe and MEA configurations. Data was collected from diverse experimental preparations across multiple species, including mole rats, monkeys, and humans. The preprocessing and spike-sorting methods were largely similar for the different Neuropixels recordings, except for the use of Kilosort 4 instead of 2.5 for spike sorting. For human recordings, the spike-sorter Combinato was used to account for noisy recording conditions in the clinic^39^. Aside from inter-species differences, these different recording and analysis conditions may have contributed to the need for re-training to achieve high classification performance in human data.

### Manual Curation

Human annotations for the base dataset were generated using a standardized manual curation procedure. The spike sorting results were curated in Phy 2.0b1^10^ by experienced curators. To ensure that curation performance was as high as possible, all curators went through repeated rounds of training and evaluated their labeling results together to ensure that classification criteria were applied consistently. Units were classified into four categories: (1) single-unit activity (SUA), representing well-isolated individual neurons; (2) multi-unit activity (MUA), representing activity from multiple neurons; (3) noise, representing non-neural artifacts (electrical, optical, or other non-biological signals); and (4) putative units, where curators were not sufficiently confident of the cluster label, for example a low-quality MUA cluster that could also be labeled as noise. To ensure clear cluster labels in the training dataset, these putative units were excluded from the initial model training.

Each dataset in the base collection was curated by at least two trained curators. Clusters labeled as putative units by the majority of curators were excluded. Agreement rates between curators were calculated separately for noise and SUA classifications, providing an upper bound on the expected classifier performance in subsequent analyses.

### Quality metric calculation and redundancy

We computed a comprehensive set of unit-level quality metrics to enable the automated cluster classification. These metrics were grouped into three categories: (1) standard quality metrics, (2) template-based metrics, and (3) newly introduced metrics. All metrics were calculated using SpikeInterface (version 102.0) in accordance with the official tutorial^11^. New quality metrics were all integrated in spike interface so that all metrics that were used in the study can be readily computed for new datasets. Detailed definitions and computation methods for each metric are provided in the appendix.

Before model training and evaluation, all feature values were converted to a 32-bit floating-point format, infinite values were replaced with `NaN`, and extreme outliers were clipped to the range [–1 × 10^6^, 1 × 10^6^] to ensure numerical stability. Log normalization was applied to spike-related features (including firing rate, SNR, and amplitude metrics) to correct for their highly skewed distributions, typical of electrophysiological data. This preprocessing pipeline was consistently applied to both training and test datasets to preserve data integrity and comparability across experimental conditions.

We examined redundancy among cluster metrics and evaluated their ability to distinguish noise, multi-unit activity (MUA), and single-unit activity (SUA) clusters. To quantify pairwise relationships between metrics, we computed the Pearson correlation matrix across all curated clusters in the base dataset (Figure 2A). Metrics were grouped into three categories: template metrics (green), quality metrics (red), and newly introduced metrics (blue). Hierarchical clustering was then applied within each category to identify groups of highly correlated metrics, which appear as distinct clusters in the correlation matrix.

To examine redundancy and shared variance among cluster metrics, we performed principal component analysis (PCA) on the full set of metrics and plotted the cumulative explained variance of the resulting principal components (Figure 2B). Prior to PCA, missing values were imputed using median substitution. All features were then standardized using the `StandardScaler()` function from scikit-learn to ensure equal contribution of all metrics with different scales (e.g., firing rate versus amplitude-based measures). The dimensionality was then determined by computing the lowest number of components required to retain 90% of the total variance, corresponding to 22 components in our dataset.

### AUC analysis for metric performance evaluation

The performance of individual quality metrics and machine learning classifiers was evaluated using the area under the receiver operating characteristic curve (AUC). For each metric and classifier, we calculated the AUC for two classification tasks: (1) distinguishing noise from non-noise (neural signals), and (2) identifying single-unit activity (SUA) from other signal types. For individual metrics, AUC values were computed by varying the decision threshold across the full range of the metric and calculating the sensitivity and specificity at each threshold. Metrics were then ranked by their median AUC values to identify the most discriminative individual features for each classification task. The AUC ranges from 0 to 1, with both 0 and 1 indicating perfect classification (for the opposing class, respectively) and 0.5 indicating random guessing. For each experimental recording from the base dataset, cluster-level quality metrics and ground truth annotations were processed independently to preserve dataset-specific distributions. Feature-level AUC-ROC scores were computed using the `roc_curve()` and `auc()` functions from scikit-learn. Each feature was treated as an independent classifier by using its raw values as prediction scores against the corresponding ground truth labels. To assess whether AUC-ROC distributions differed from chance, we used bidirectional Wilcoxon signed-rank tests for each metric.

### Machine Learning Methodology in UnitRefine

UnitRefine consists of a comprehensive supervised machine learning framework for automated spike sorting curation. The framework consists of two complementary components: model training and model deployment, organized around two core modules:

**Training pipeline** (train_manual_curation.py): enables researchers to train new custom models on their own manually curated datasets.

**Model based curation pipeline** (model_based_curation.py): provides application of pretrained models to new spike sorting outputs.

Both modules can be used programmatically in Python or through the UnitRefine GUI. UnitRefine runs within the SpikeInterface ecosystem, and no additional software packages are required beyond the SpikeInterface and its standard dependencies.

#### Training Pipeline

The pipeline develops custom models through the `CurationModelTrainer` class, automating all major steps of model development. The trainer accepts expert-curated human labels and spike sorting quality metrics, and then selects data preprocessing, model selection methodology, and performance evaluation. Users can specify the set of features (metrics) to include, define pre-processing options such as missing-value imputation, feature scaling, select among multiple machine learning algorithms, and apply optional data balancing for imbalanced datasets (which is often the case for SUA classification). Hyperparameter optimization is handled automatically using Bayesian search methods. To accommodate different experimental workflows, UnitRefine supports two flexible data input modes:

1. Analyzer-based training integrates directly with `SortingAnalyzer` objects in SpikeInterface to compute and extract cluster metrics and ensure parameter consistency across recordings. This is ideal for using UnitRefine as an automatic curation step as part of a spike-sorting pipeline in SpikeInterface.
2. File-Based Training allows the training and application of models based on pre-computed quality metrics and labels that are provided in comma-separated value (.csv) files. This allows flexible integration of UnitRefine in custom workflows that do not require SpikeInterface but aim to use UnitRefine as a stand-alone curation tool.

For classification, UnitRefine supports a diverse ensemble of algorithms, including tree-based ensembles (Random Forest, AdaBoost, Gradient Boosting), advanced gradient boosting frameworks (XG-Boost, LightGBM, CatBoost), and linear or neural baselines (SVC, Logistic Regression, MLP). All classifiers can be initialized with fixed random seeds to ensure reproducibility.

Model hyperparameters are tuned using an automated search framework. The primary optimization strategy employs Bayesian optimization to efficiently explore the parameter space, with randomized search available as a fallback for broad compatibility across algorithms. All model optimization relied on five-fold stratified cross-validation. Balanced accuracy was used as the evaluation metric to mitigate class imbalance. We report 95% confidence intervals derived from the empirical distribution of balanced accuracy across cross-validation folds, using the 2.5th and 97.5th percentile boundaries.

The example below shows how to train a UnitRefine model using the default Random Forest classifier on a labeled dataset. The same interface can be used with either SpikeInterface `SortingAnalyzers` or pre-computed csv files. Depending on the mode, either the `analyzers` variable for the object or the `metrics_path` variable for the .csv file need to also be specified.

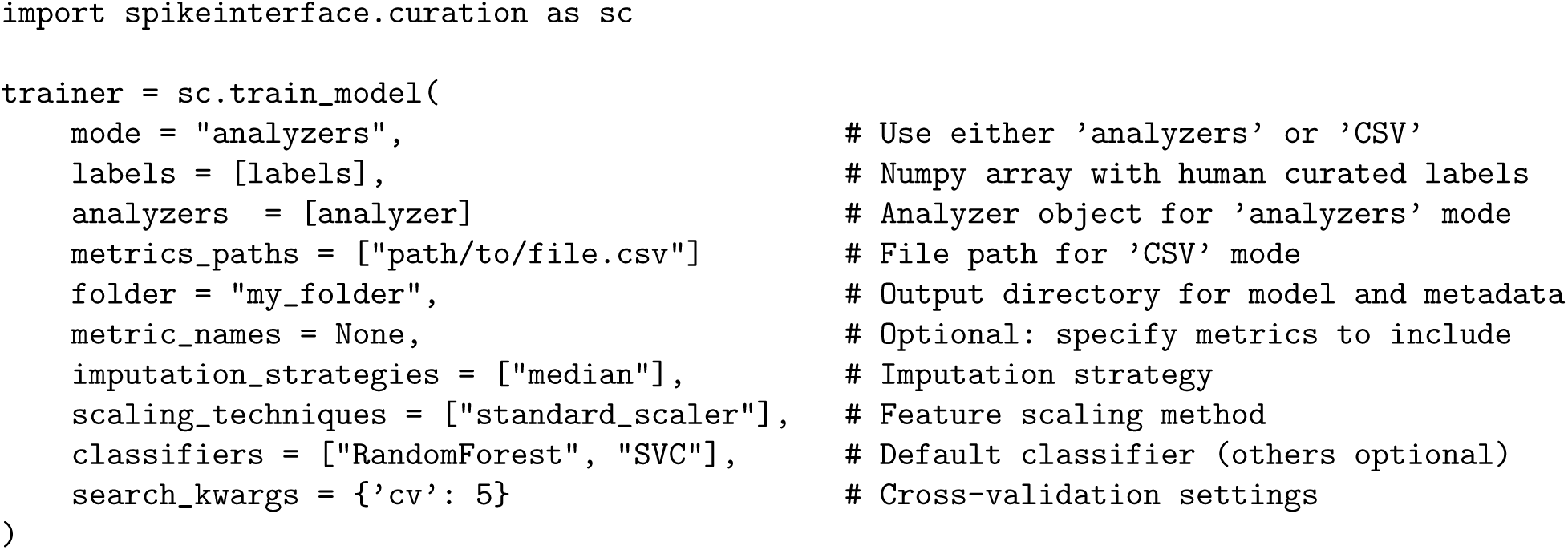

This command runs the complete UnitRefine training pipeline, including data preprocessing, cross-validation, and model selection. The best-performing classifier and its configuration are automatically saved to the specified folder in two files: `model.skops` and `model_info.json`. The `skops` file uses a secure serialization format for scikit-learn models, preserving all training and preprocessing parameters (e.g., imputation, scaling, feature order, fitted model). The accompanying `model_info.json` file records the features used during training, label mappings, class names, and software dependencies to ensure full reproducibility.

The trained UnitRefine models can be shared on the Hugging Face Hub for community reuse and benchmarking. Each uploaded model includes `model.skops` and `model_info.json` files. An additional `metadata.json` file records essential biological and acquisition information (e.g., species, brain areas, probe type) to guide appropriate reuse. The metadata fields are formatted to align with the Neurodata Without Borders (NWB) conventions^40^. A metadata example for a mole rat recording is organized as follows:

~~~
model_metadata = {
   “subject_species”: [“Mus_musculus”],
   “brain_areas”: [“V1”],
   “probes”: [{
     “manufacturer”: “IMEC”,
     “name”: “Neuropixels 2.0”
   }]
}
~~~

Once the metadata file is created, the entire model directory (`my_folder`) can be uploaded directly to the Hugging Face Hub, enabling model sharing, comparison, and reuse across laboratories. A typical model folder has the following structure:

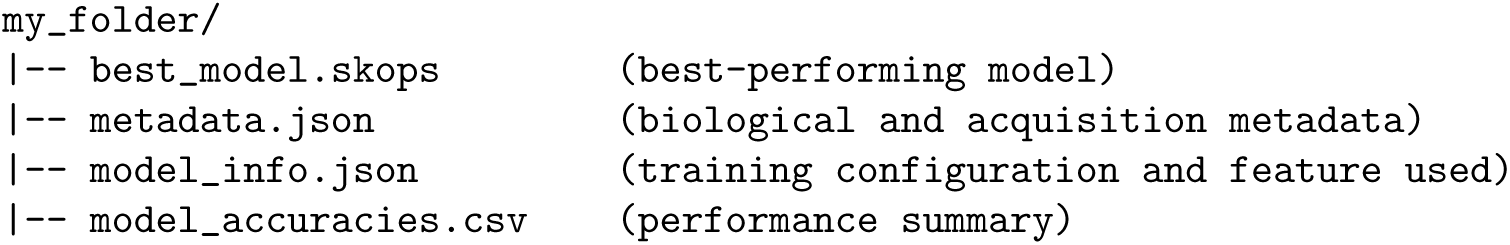
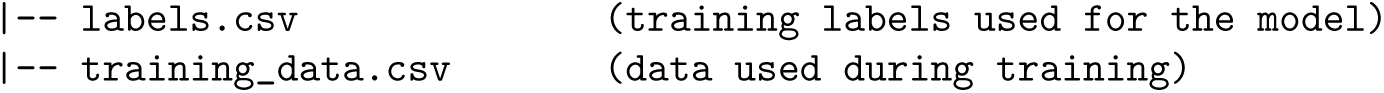

#### Model-based curation pipeline

This module allows users to apply a pretrained machine learning model to curate their own spike-sorted data. When a trained model directory (`my_folder`) is uploaded to the Hugging Face Hub, its contents form the pretrained model package used by the model-based curation pipeline. Within this package, `best_model.skops` is the file that UnitRefine loads to generate predictions on new spike-sorted data.

The `ModelBasedClassification` class is designed with the following capabilities: 1) Flexible model loading: supports loading models either from a local directory or directly from the Hugging Face Hub. 2) Parameter and feature validation: ensures that all required metrics are available, feature names match, and preprocessing parameters are consistent with those used during training. 3) Confidence reporting: outputs predicted labels along with calibrated probabilities for each cluster to support downstream quality control.

The example below shows how to retrieve a public model from Hugging Face Hub and apply it to a `SortingAnalyzer` object. The output is a pandas dataframe with the predicted labels. The same functions also accept a local `folder` path instead of `repo_id`.

~~~
# Download a pretrained model (or load from a local folder).
model, model_info = sc.load_model(
   repo_id = “SpikeInterface/UnitRefine_sua_mua_classifier”,
   trusted = [“numpy.dtype”]  # whitelist for secure deserialization
)
~~~

~~~
# Predict labels for units in a SortingAnalyzer.
labels = sc.auto_label_units(
   sorting_analyzer = sorting_analyzer,
   repo_id = “SpikeInterface/UnitRefine_sua_mua_classifier”,
   trusted = [“numpy.dtype”]
)
~~~

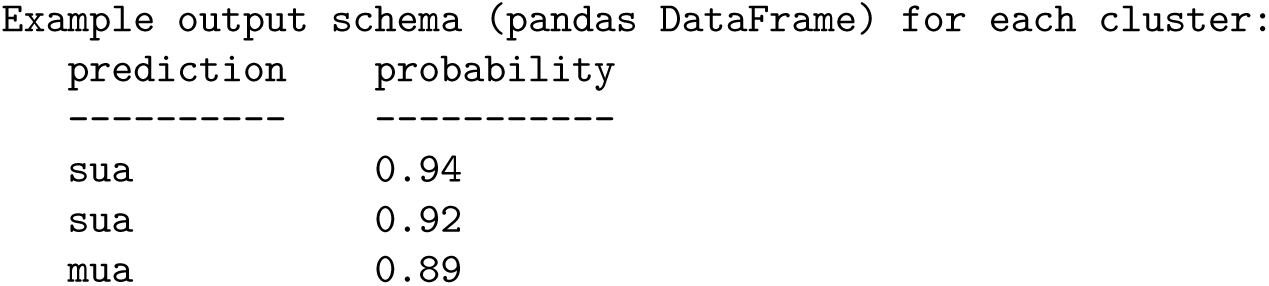

| prediction | probability |
| --- | --- |
| sua | 0.94 |
| sua | 0.92 |
| mua | 0.89 |
| ... |  |

To obtain an estimate of the computation time for each classifier with different metric sets, we analyzed a 25-minute-long NeuroPixels 1.0 recording with 384 channels and 242 units. The preprocessed recording was then saved locally before computing all the required extensions. The computation was done on an 8-core workstation with an Intel i-7 CPU and 32 GB of RAM.

### SHAP-Based Model Interpretation

To interpret how individual quality metrics influenced classification decisions, we applied SHapley Additive exPlanations (SHAP) analysis. SHAP decomposes each model prediction into additive contributions from all input features, indicating how much each metric increases or decreases the probability of an individual cluster being classified as SUA and providing a consistent and interpretable visualization of the decision process.

For each dataset and cross-validation fold, SHAP feature importances (Figure 4B, F) were computed using the `TreeExplainer` algorithm, which is specifically optimized for tree-based models such as Random Forest decoders. The SHAP feature importance workflow comprised three main steps: 1) During model training, Random Forest classifiers were trained for each random seed (*n* = 10) on pre-processed data. 2) After training, the Random Forest estimator was extracted from the `scikit-learn` pipeline, and SHAP values were computed for the test set of each fold, focusing on the positive SUA class. 3) Lastly, feature importance was quantified as the mean absolute SHAP value across all test samples within a fold, providing a measure of the average contribution of each feature to the model’s predictions.

Feature importance was visualized as sorted bar plots displaying mean absolute SHAP values, with error bars representing the standard deviation across random seeds. For cross-species and cross-dataset comparisons (Figure 4K and Figure 5F), SHAP-based feature rankings were visualized as heatmaps, highlighting differences in feature importance across species (mouse, mole rat, monkey, human) and datasets. Lower rank values corresponded to higher importance (i.e. the first feature being the most important), facilitating direct comparison across experimental contexts.

To further investigate model predictions, complementary SHAP summary and decision plots were used to examine how variations in individual feature values affected the model’s output probabilities. Summary plots provide an overview of feature importance distributions across all clusters, while decision plots show the cumulative feature contributions along individual prediction paths, therefore visualizing model decisions at both the population and single-unit levels.

### Decoding of behavior in a decision-making task

We reproduced the methodology by using the dataset and code described in the IBL brain-wide-map analysis^6^ to decode two task variables: feedback outcome and choice. To compute decoder accuracy, we used validated maximum-likelihood regression with L1 regularization. Cross-validation used a null distribution to test for significance. To aggregate neurons from the same region but multiple recordings, we combined P values using Fisher’s combined probability test.

We performed three decoding runs with the setup above. First we reproduced the results from the main paper using cluster metric thresholds to select the units. Second, we used the Unit-Refine model trained on data acquired under the same task^29^. In the third and last run, to control for the effect of the number of units, we selected all of the threshold units from the original paper, and randomly selected units from the remaining pool until we reached the same unit count that Unit-Refine yielded.

### GUI and Active learning

To facilitate accessibility for experimental laboratories and non-programmatic users, we developed a graphical user interface (GUI) for UnitRefine using the `PyQt5` framework. The GUI provides a guided workflow for model training and automated curation, organized into four main steps (Figure 6F).

In the first step, users create a project directory. In the second step they load spike-sorting outputs as a `SortingAnalyzer` object. Multiple analyzer objects can be imported simultaneously from either local storage or cloud sources. Selecting an uploaded analyzer opens the integrated SpikeInterface-GUI visualization backend, which displays waveforms, spike amplitudes, correlograms, and firing rates. Users can then add or correct labels using hotkeys, and assigned labels appear in the `quality` column (Supplementary Figure S5).

The third step provides an interface for training new models or loading existing UnitRefine models from disk or directly from the Hugging Face Hub. The training menu (Supplementary Figure S6) shows all configurable components, including metric selection, classifiers, scalers, imputers, and test-size parameters. Predicted labels are displayed for validation through Spikeinterface-GUI and users can provide additional labels.

The final step introduces an integrated active learning workflow (Figure 6G), implemented using the `modAL` library^41^. After an initial model is trained, clusters with low prediction confidence are automatically identified and presented to the user for targeted relabeling. These updated labels are incorporated into the training set, enabling iterative retraining and progressive improvement of model performance. Completed model folders can be uploaded to the Hugging Face Hub for distribution and reuse across laboratories.

## Acknowledgements

CH and MHH are supported by the UKRI Biotechnology and Biological Sciences Research Council (BBSRC) grant number BB/X01861X/1. RG is supported by the UKRI Biotechnology and Biological Sciences Research Council (BBSRC) grant number BB/T00875X/1.

SM is supported by the Helmholtz Association VH-NG-1611. AJ, SM, and SG are supported by the state of North Rhine-Westphalia through the iBehave initiative (grant number NW21-049).

SG is funded by the Deutsche Forschungsgemeinschaft (DFG, German Research Foundation) - 368482240/GRK2416, Horizon Europe Grant 101147319, and DFG grant 561027837/ GR 1753/9-1.

We express our gratitude to our curators: Nilufar Lahiji, Sacha Abou Rachid, Severin Graff, Luca Koenig, and Natalia Babushkina for their invaluable help in curating the data.

The authors also thank Alana Darcher and Dr. Florian Mormann for discussions and providing the human patient data, Aitor Morales-Gregorio for contributing macaque data, Jake Swann for sharing rat data, Sylvia Schroeder for Neuropixels data, and Alireza Saeedi, Runita Shirshankar, and Pascal Malkemper for providing data from Fukomys anselli. We also thank Joe Ziminski for giving valuable feedback during the SpikeInterface integration process.

## Appendix: Quality Metric Descriptions

We exclusively used the implementations of quality and template metrics contained in the SpikeInter-face package^10^ for model training. These are re-implementations of metrics from a diverse range of papers and from GitHub repositories. We have tried to cite the first appearance of each metric in the literature, though this is difficult for very commonly used metrics.

The `firing_range`, `amplitude_cv_median`, `amplitude_cv_range`, and `sync_n` metrics are newly developed metrics that we integrated into SpikeInterface. Below, we provide an overview of the metric definitions using unified notation. All metrics listed here were computed for each cluster and used in the full decoder models.

### Terminology, conventions and notation

Cluster: Set of spikes, determined by a spike sorting algorithm to be clustered together.

Waveform: The preprocessed voltage trace of a spike. Can exist on many probe channels.

Template: The representative waveform of a single cluster, usually computed as the mean or median of the cluster’s spike waveforms.

We index spikes with lower case letters, and clusters with upper case letters.

*T* : total time of the recording (*s*)

*f* : sampling frequeny (*Hz*)

*N* : number of spikes

*N_I_* : number of spikes in cluster *I*.

### Spike train metrics

`num_spikes`

Number of spikes in the cluster.

`firing_rate`

Average number of spikes in the cluster, per second.

`firing_range`

For each cluster, the spike train is binned in time and a firing rate is computed for each time bin. This produces a distribution of firing rates. The firing range is the difference between the 5th and 95th percentile firing rates across the recording.

`presence_ratio`

For each cluster, the spike train is binned in time. The presence ratio is equal to the fraction of bins containing at least one spike.

`sync_n`

Take all spike times from cluster *I*. Count all spikes which occur at the same timepoint (sample) as at least *n* − 1 spikes from all other clusters. `sync_n` is equal to this count.

### Refractory period violation metrics

#### Additional notation

*T_r_* : refractory period (s).

Once a neuron has fired an action potential, it cannot do so again for a duration known as the refractory period. If two spiking events occur within this time interval, a refractory period violation has occurred. The length of the refractory period is different in different neuron types, but usually lasts for a few milliseconds.

`isi_violations_count`

The number of inter-spike-interval (isi) violations that occured for each cluster. Here, the number of violated refractory periods are counted. Hence if two spikes violate one period, only one violation is counted (in contrast with `rp_violations`).

isi_violations_ratio^15^

Given the `isi_violations_count`, the `isi_violations_ratio` approximates the contamination rate of the cluster, assuming random contamination. For a cluster *I*, the approximation is given by

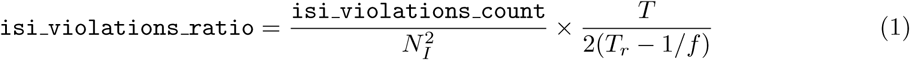

`rp_violations`

The number of refractory period (rp) violations that occur for each cluster. Here the total number of violations is counted. Hence if two spikes violate another spikes’ refractory period, two violations are counted (in contrast with `isi_violations`).

`rp_contamination`^42^

Given the `rp_violations` count, the contamination rate of the cluster is approximated, assuming random contamination. The approximation is a refinement of `isi_violations_ratio` and works better for highly contaminated clusters. For a given cluster *I*, the ratio is given by

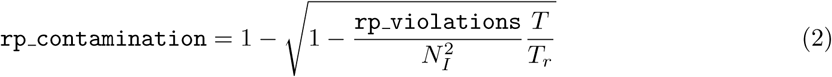

`sliding_rp_violation`^30^

Assume that a percentage of the cluster’s spikes come from contamination. Compute the likelihood of observing the computed `isi_violations` for a given refractory period, assuming Poisson spiking. Do this for several putative refractory periods. If the confidence is above a 90% confidence threshold, keep this refractory period. The `sliding_rp_violation` is equal to the minimum contamination rate from all kept refractory periods. Allows a metric to be computed for neurons with an unknown refractory period.

### Amplitude metrics

#### Additional notation

spike amplitudes: Using either `max`, `min` or `max abs` as the extremal function, find the channel whose cluster template has the maximal extremal value. Call this the extremal channel. For each spike, the spike amplitude is equal to the extremal amplitude on the extremal channel.

amplitude cv: Consider the distribution of spike amplitudes for some subset of spikes. The amplitude coefficient of variation (cv) is equal to the standard deviation of this distribution, divided by the mean.

`snr`

The snr is equal to the extremal amplitude of the template on the extremal channel, divided by the standard deviation of the noise on this channel.

`sd_ratio`^43^

Compute the spike amplitudes for a simulated cluster composed of many snippets of noise, giving a noise-amplitude distribution. The `sd_ratio` is equal to the standard deviation of the cluster’s amplitude distribution divided by the standard deviation of the noise-amplitude distribution.

`amplitude_cv_median`

For each cluster, time bin the spike amplitudes and compute the `amplitude_cv` for each bin. The `amplitude_cv_median` is the median of the values across all bins in a given recording.

`amplitude_cv_range`

For each cluster, time bin the spike amplitudes and compute the `amplitude_cv` for each bin. Find the 5^th^ and 95^th^ percentile values. The `amplitude_cv_range` is the difference between these values.

`amplitude_median`

The median of the distribution of the spike amplitudes for each cluster.

`amplitude_cutoff`^5^

For each cluster, make a histogram of spike amplitude values. Assume the distribution is normally distributed and fit the distribution using the high-value tail. Under this assumption, estimate how many spikes are missing from the low-value tail. The `amplitude_cutoff` is equal to the estimated fraction of spikes missing from the tail.

### Drift metrics

#### Additional notation

Spike location difference distribution: for each cluster, compute the location of each spike, giving a distribution of spike locations relative to the probe. Then compute the difference between the locations and the median location. This gives a distribution of spike location differences for each cluster.

`drift_ptp`

The difference between the maximum and minimum values in the spike location difference distribution.

`drift_std`

The standard deviation of the spike location difference distribution.

`drift_mad`^5^

The median absolute deviation of the spike location difference distribution. More precisely,

~~~
drift_mad = median (abs (spike_location_difference − mean (spike_location_difference))).
~~~

### Principal Components metrics

For these metrics, the principal components of each spike waveform are computed. When distance is used, this refers to the distance measured by a metric, most commonly the *L*_2_ metric, in principal component space. The Mahalanobis distance accounts for cluster means and variations.

`isolation_distance`^44^

Compute the the Mahalanobis distance between cluster *I* and all other clusters. The `isolation_distance` for cluster *I* is equal to the minimum of all these distances.

`silhouette`^45^

Take a cluster I. For every spike *i* ∈ *I*, compute the average distance to every other spike from its own cluster and call this *a*(*i*). Call the other clusters *B*_1_*, B*_2_, Now compute the average distance from *i* to every spike in each of these clusters and call these distances *b*_1_(*i*)*, b*_2_(*i*), Take the shortest of these distances, *b*(*i*) = min*_I_ b_I_* (*i*). The sillhoutte score of this spike is equal to

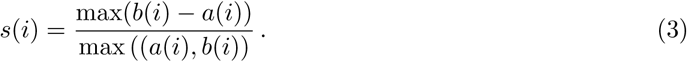

The cluster silhouette score of the cluster, silhouette, is the average of s(i) across all spikes.

`nn_hit_rate`^5^

For each spike in a cluster *A*, find `n` nearest neighbour spikes. The spike hit rate is the fraction of these spikes which are also in *A*. The cluster spike rate is the mean of all spike hit rates for spikes in *A*.

`nn_miss_rate`^5^

For each spike not in a cluster *A*, find the `n` nearest neighbour spikes. The spike miss rate is the fraction of these spikes which are in A. The cluster spike rate is the mean of all spike hit rates for spikes not in A.

`d_prime`^15^

Consider the spikes in one cluster *A*, and the set of all other spikes, ¬*A*. Construct a one-component Linear Discrimiation Model with respect to these sets, which contains a mean and standard deviation for each set of spikes *µ_A_, µ_¬A_, σ_A_, σ_¬A_*. The `d_prime` is equal to the normalised difference in these means:

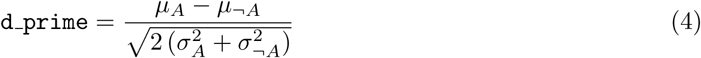

`l_ratio`^46^

Assume all principal components for each cluster come from indepdendent, normal distributions. This gives a probability *χ*^2^ distribution in PCA space. Consider all spikes that do not belong to cluster I, *s_j_*. Using the Mahalanobis distance in PCA space between the spike *s_j_* and the cluster centroid *C_I_* , we can compute the probability that the spike *s_j_* should have been sorted into *I*. It is given by

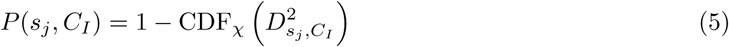

The l_ratio for a cluster *I* is then equal to

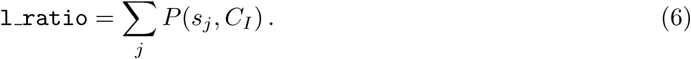

### Template metrics

The template metrics are mostly derived from Siegle et al.^5^.

#### Additional notation

A template is defined on a number of channels. The template on each channel has a `peak` (the maximum value) and a `valley` (the minimum value). We call the channel with the extremal value (either with respect to `max`, `min` or `abs_max`) the extremal channel. Many metrics (`peak_to_valley, get_peak_trough_ratio, half_width, repolarization_slope, recovery_slope, num_positive_peaks, num_negative_peaks`) only use the template from the extremal channel. We show a visual representation of several of these metrics in Figure 7.

**Figure 7:**
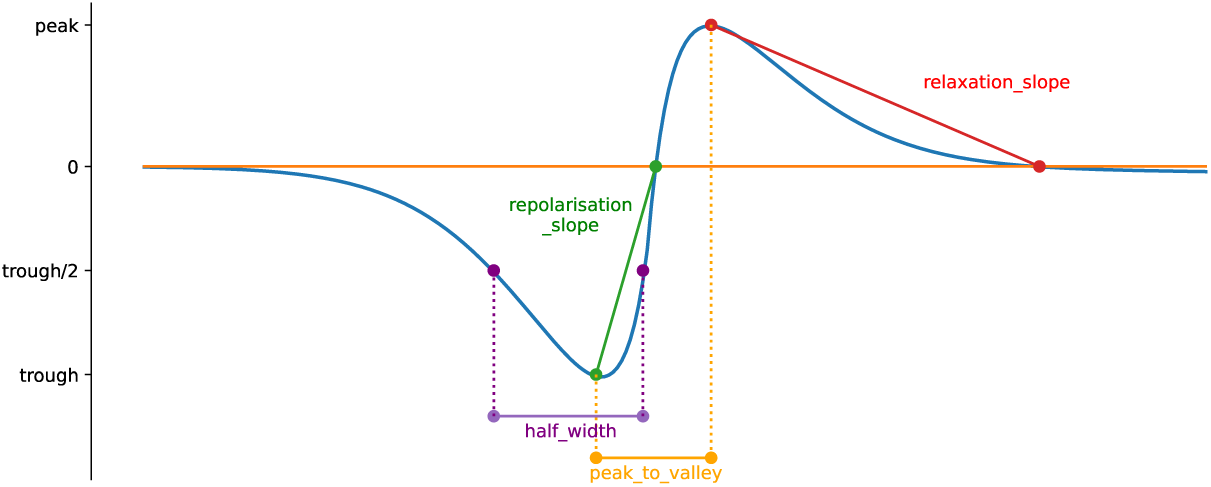
A visual representation of some of the template metrics.

`peak_to_valley (s)`

The time between the peak and valley of the template on the extremal channel.

`peak_trough_ratio`

The ratio when dividing the peak by the trough 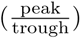.

`num_positive_peaks`

The number of positive peaks of the template on the extremal channel.

`num_negative_peaks`

The number of negative peaks of the template on the extremal channel.

`half_width (s)`

Consider the `trough`. The `half_width` is equal to the maximum interval around the valley index where all template values lie between the `trough` value and half of the `trough` value.

Repolarization_slope (*µV/s*)

Consider the line from the `trough` to the first zero-value following the `trough`. The `repolarization_slope` is the gradient of this line.

recovery_slope (*µV/s*)

Consider the line from the `peak` to the first zero-value following the `peak`. The `recovery_slope` is the gradient of this line.

exp_decay (1*/µm*)

On every channel, find both the maximum amplitude and the distance from the extremal channel. Use this to produce amplitude as a function of distance *A*(*d*). Fit this function to an exponential decay *A*(*d*) ≈ *A*_0_ exp(−*ad*). The metric exp_decay is equal to the decay rate *a*.

`velocity_above`

On every channel above the extremal channel, find the time index of the extremal value of the template and the distance from the extremal channel. Using this, produce distance as a function of extremal-amplitude-temporal-location, *d*(*t*). Fit this relation using a linear regression. The gradient represents the velocity of the spike, which equals `velocity_above`.

`velocity_below`

On every channel below the extremal channel, find the time index of the extremal value of the template and the distance from the extremal channel. Using this, produce distance as a function of extremal-amplitude-temporal-location, *d*(*t*). Fit this relation using a linear regression. The gradient represents the velocity of the spike, which equals `velocit_below`.

`spread`

Find all channels containing a channel-template whose amplitude is above a given threshold. The `spread` is equal to the maximum distance between the channels that pass this threshold.

## Supplementary Tables

**Table 1:** Results from different classification algorithms for decoding noise versus neural clusters.

| Classifier | Median (%) | Mean (%) | 95% Confidence interval |
| --- | --- | --- | --- |
| RandomForest (RF) | 84.2 | 83.0 | 73.8 – 90.9 |
| LGBMClassifier (LGBM) | 82.8 | 80.5 | 68.0 – 89.3 |
| GradientBoosting (GB) | 84.3 | 79.7 | 65.7 – 90.0 |
| XGBClassifier (XGB) | 82.7 | 78.4 | 66.1 – 89.8 |
| AdaBoost (Ada) | 79.4 | 78.0 | 67.4 – 90.3 |
| TabPFN | 80.3 | 77.8 | 65.5 – 90.1 |
| Bombcell | 78.0 | 77.4 | 64.7 – 89.9 |
| LogisticRegression (LR) | 77.7 | 74.0 | 64.8 – 80.4 |
| MLP | 72.6 | 72.5 | 64.2 – 84.0 |

**Table 2:**
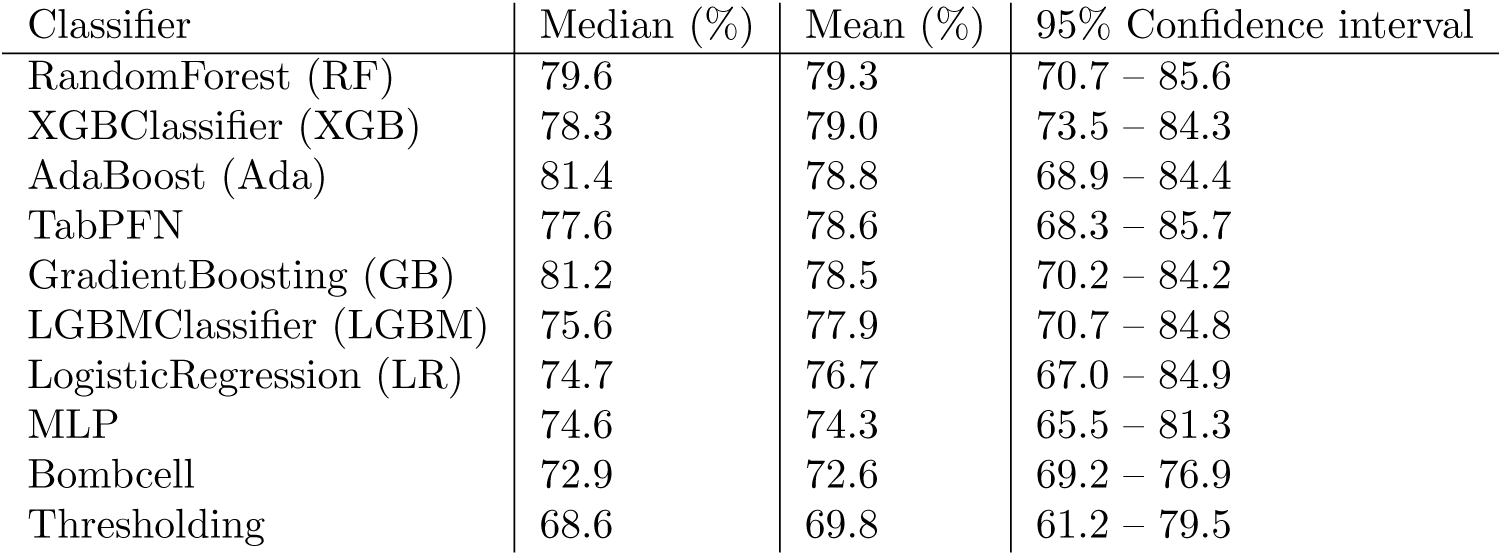
Results from different classification algorithms for decoding MUA versus SUA clusters.

| Classifier | Median (%) | Mean (%) | 95% Confidence interval |
| --- | --- | --- | --- |
| RandomForest (RF) | 79.6 | 79.3 | 70.7 – 85.6 |
| XGBClassifier (XGB) | 78.3 | 79.0 | 73.5 – 84.3 |
| AdaBoost (Ada) | 81.4 | 78.8 | 68.9 – 84.4 |
| TabPFN | 77.6 | 78.6 | 68.3 – 85.7 |
| GradientBoosting (GB) | 81.2 | 78.5 | 70.2 – 84.2 |
| LGBMClassifier (LGBM) | 75.6 | 77.9 | 70.7 – 84.8 |
| LogisticRegression (LR) | 74.7 | 76.7 | 67.0 – 84.9 |
| MLP | 74.6 | 74.3 | 65.5 – 81.3 |
| Bombcell | 72.9 | 72.6 | 69.2 – 76.9 |
| Thresholding | 68.6 | 69.8 | 61.2 – 79.5 |

**Table 3:** Overview of the different metric groups that were included in the reduced classifiers. A detailed description of all cluster metrics is given in section .

| Metric List | Features |
| --- | --- |
| **All Metrics** | num_spikes, firing_rate, presence_ratio, snr, isi_violations_ratio, isi_violations_count, rp_contamination, rp_violations, sliding_rp_violation, amplitude_cutoff, amplitude_median, amplitude_cv_median, amplitude_cv_range, sync_spike_2, sync_spike_4, sync_spike_8, firing_range, drift_ptp, drift_std, drift_mad, isolation_distance, l_ratio, d_prime, silhouette, nn_hit_rate, nn_miss_rate, peak_to_valley, peak_trough_ratio, half_width, repolarization_slope, recovery_slope, num_positive_peaks, num_negative_peaks, velocity_above, velocity_below, exp_decay, spread |
| **Quality Metrics** | num_spikes, firing_rate, presence_ratio, snr, isi_violations_ratio, isi_violations_count, rp_contamination, rp_violations, sliding_rp_violation, amplitude_cutoff, amplitude_median, amplitude_cv_median, amplitude_cv_range, sync_spike_2, sync_spike_4, sync_spike_8, firing_range, drift_ptp, drift_std, drift_mad, isolation_distance, l_ratio, d_prime, silhouette, nn_hit_rate, nn_miss_rate |
| **Template Metrics** | peak_to_valley, peak_trough_ratio, half_width, repolarization_slope, recovery_slope, num_positive_peaks, num_negative_peaks, velocity_above, velocity_below, exp_decay, spread |
| **Spike Times Metrics** | num_spikes, firing_rate, presence_ratio, snr, isi_violations_ratio, isi_violations_count, rp_contamination, rp_violations, sliding_rp_violation, sync_spike_2, sync_spike_4, sync_spike_8, firing_range |

**Table 4:** Overview of datasets used in this study. All model performance was evaluated using a leave-one-session-out validation method.

| Dataset | Probe type | n recordings | Spike sorter | Species |
| --- | --- | --- | --- | --- |
| Base dataset | Neuropixels 1.0 | 11 | Kilosort 2.5 | Mouse |
| IBL | Neuropixels 1.0 | 8 | IBL-sorter (PyKS2.5) | Mouse |
| Allen Institute for Neural Dynamics | Neuropixels 2.0 | 4 | Kilosort 4 | Mouse |
| Mole rat recordings | Neuropixels 2.0 | 4 | Kilosort 4 | Mole rat |
| Nonhuman primate recordings | Utah array | 11 | Kilosort 4 | Macaque |
| Human intracranial recordings | Behnke–Fried electrodes | 12 | Combinato<sup>39</sup> | Human |

**Table 5:**
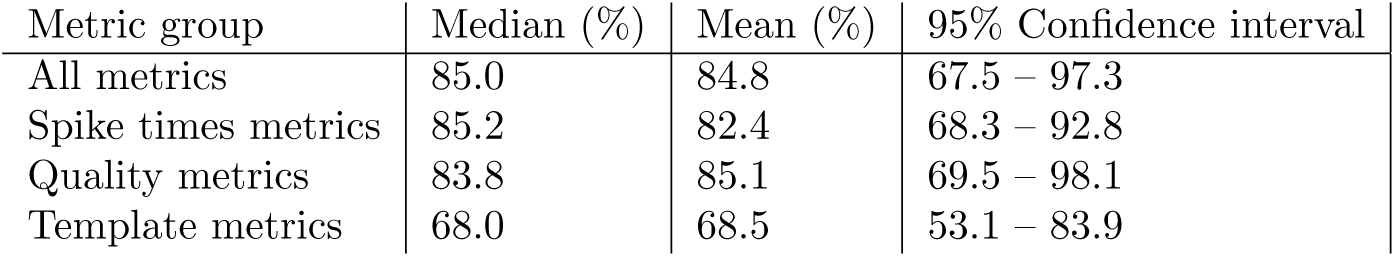
Median, mean, and 95% confidence interval of balanced accuracy performance across all tested recordings for different metric groups.

| Metric group | Median (%) | Mean (%) | 95% Confidence interval |
| --- | --- | --- | --- |
| All metrics | 85.0 | 84.8 | 67.5 – 97.3 |
| Spike times metrics | 85.2 | 82.4 | 68.3 – 92.8 |
| Quality metrics | 83.8 | 85.1 | 69.5 – 98.1 |
| Template metrics | 68.0 | 68.5 | 53.1 – 83.9 |

**Table 6:**
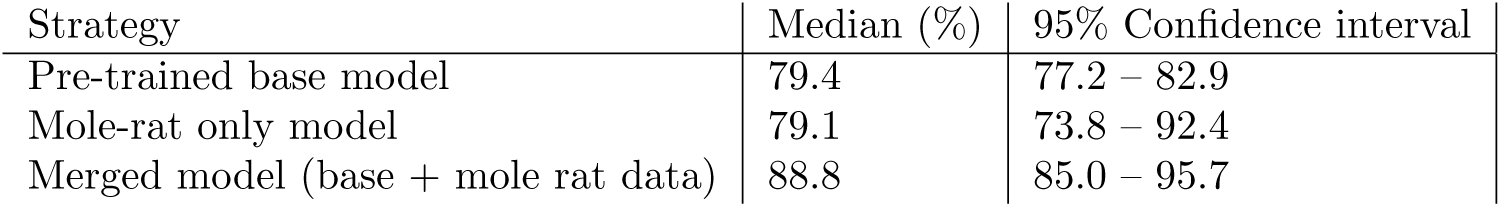
Median and 95% confidence interval for different training strategies for curation of molerat data.

| Strategy | Median (%) | 95% Confidence interval |
| --- | --- | --- |
| Pre-trained base model | 79.4 | 77.2 – 82.9 |
| Mole-rat only model | 79.1 | 73.8 – 92.4 |
| Merged model (base + mole rat data) | 88.8 | 85.0 – 95.7 |

**Table 7:** Median and 95% confidence interval for different training strategies for curation of monkey data.

| Strategy | Median (%) | 95% Confidence interval |
| --- | --- | --- |
| Pre-trained base model | 73.7 | 56.2 – 96.7 |
| Monkey only model | 93.1 | 76.9 – 99.0 |
| Merged model (base + monkey data) | 91.7 | 86.0 – 99.8 |

**Table 8:** Median and 95% confidence interval for different training strategies for curation of human data.

| Strategy | Median (%) | 95% Confidence interval |
| --- | --- | --- |
| Pre-trained base model | 39.6 | 23.8 – 54.2 |
| Human only model | 77.5 | 68.4 – 87.9 |
| Merged model (base + human data) | 74.2 | 69.8 – 89.0 |

## Supplementary Figures

**Figure S1:**
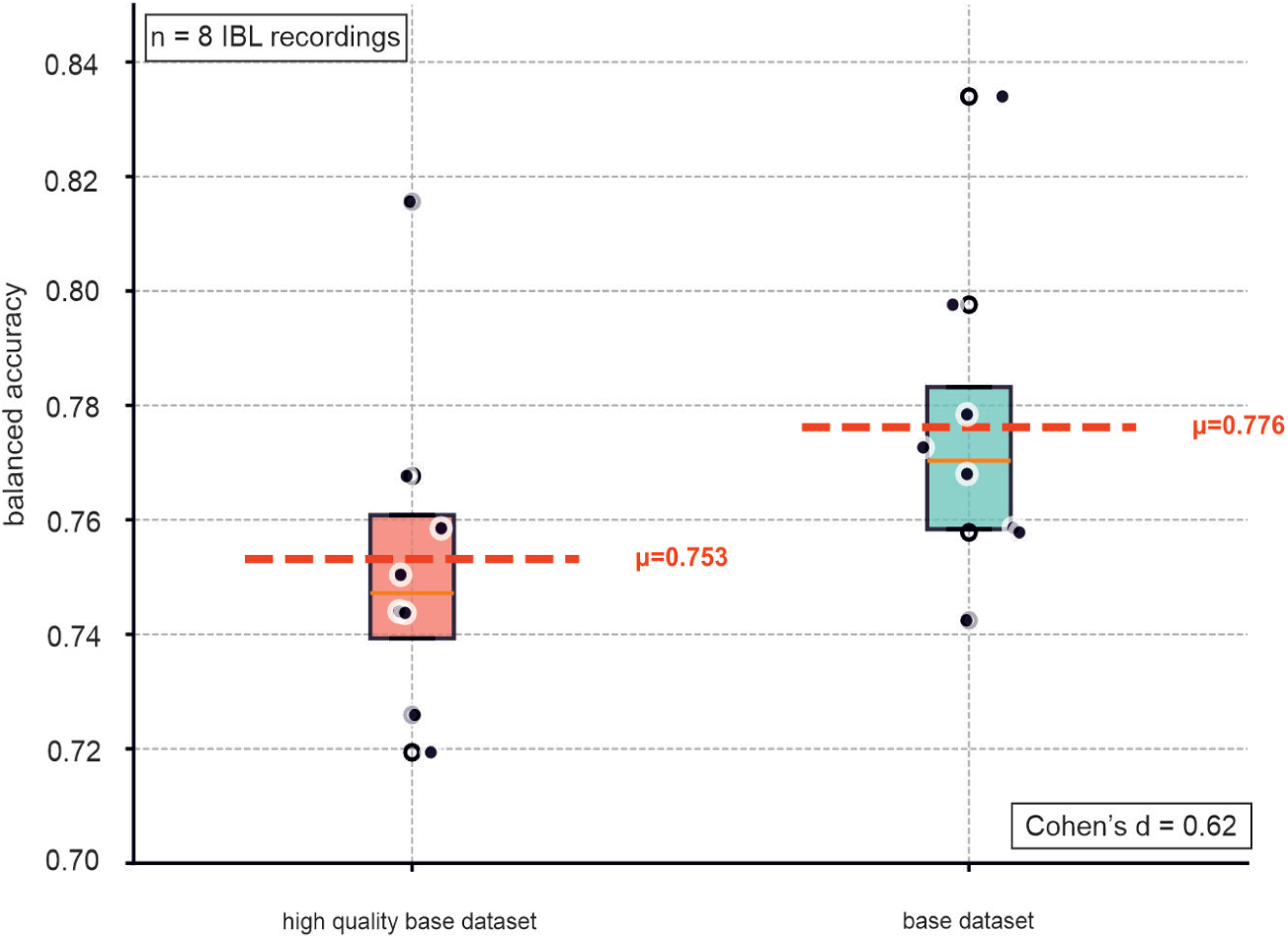
Performance comparison between high-quality and base dataset models. Balanced accuracy scores are shown for two models, either trained on the high-quality or the full base dataset on *n* = 8 IBL recordings. The base-traine model achieved slightly higher average performance (*µ* = 0.776) compared to the high-quality-trained model (*µ* = 0.753). The moderate effect size (Cohen’s *d* = 0.62) indicates a consistent improvement when incorporating a broader diversity of training data.

**Figure S2:**
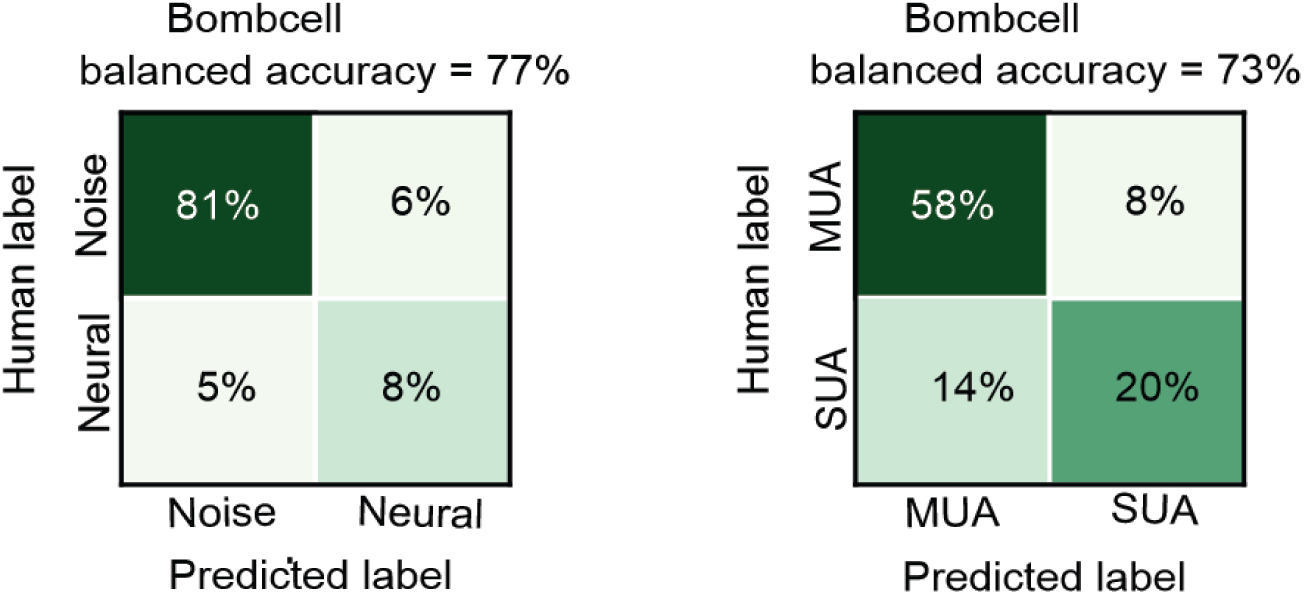
Performance of Bombcell classifier. (**A**) Confusion matrix showing the performance of the noise vs. neural classifier, achieving a balanced accuracy of 77%. (**B**) Confusion matrix showing the performance of the MUA vs. SUA classifier, achieving a balanced accuracy of 75%.

**Figure S3:**
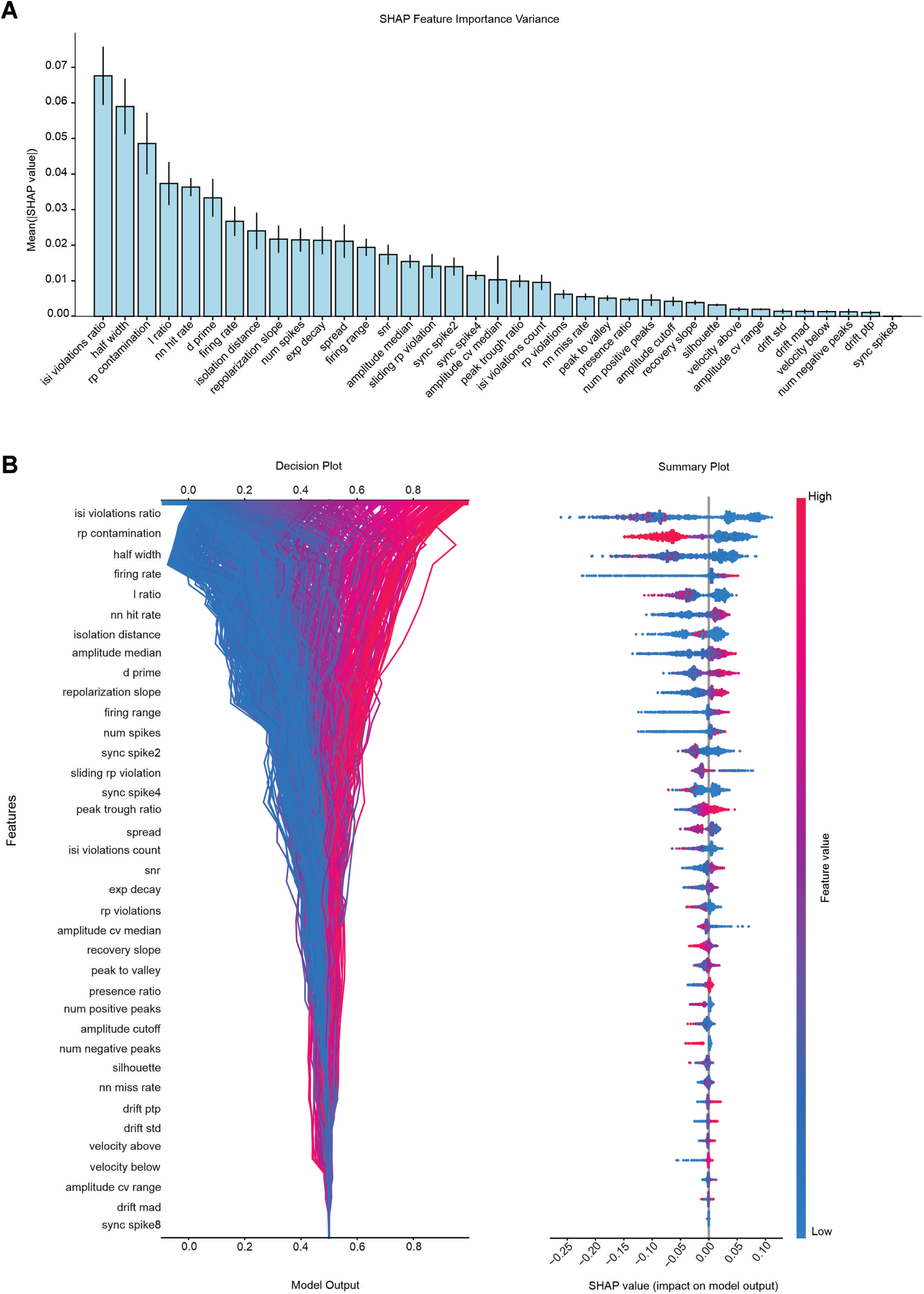
SHAP-based interpretation of UnitRefine classifier decisions. (**A**) Mean SHAP feature importance values with standard deviation across ten cross-validation folds, showing the relative contribution of each quality metric to model predictions. Same das as in Figure 3H but using all available cluster metrics. Most metrics made signfiicant contributions to decoder performance. (**B**) SHAP decision and summary plots illustrating how all features influence the model output for individual clusters.

**Figure S4:**
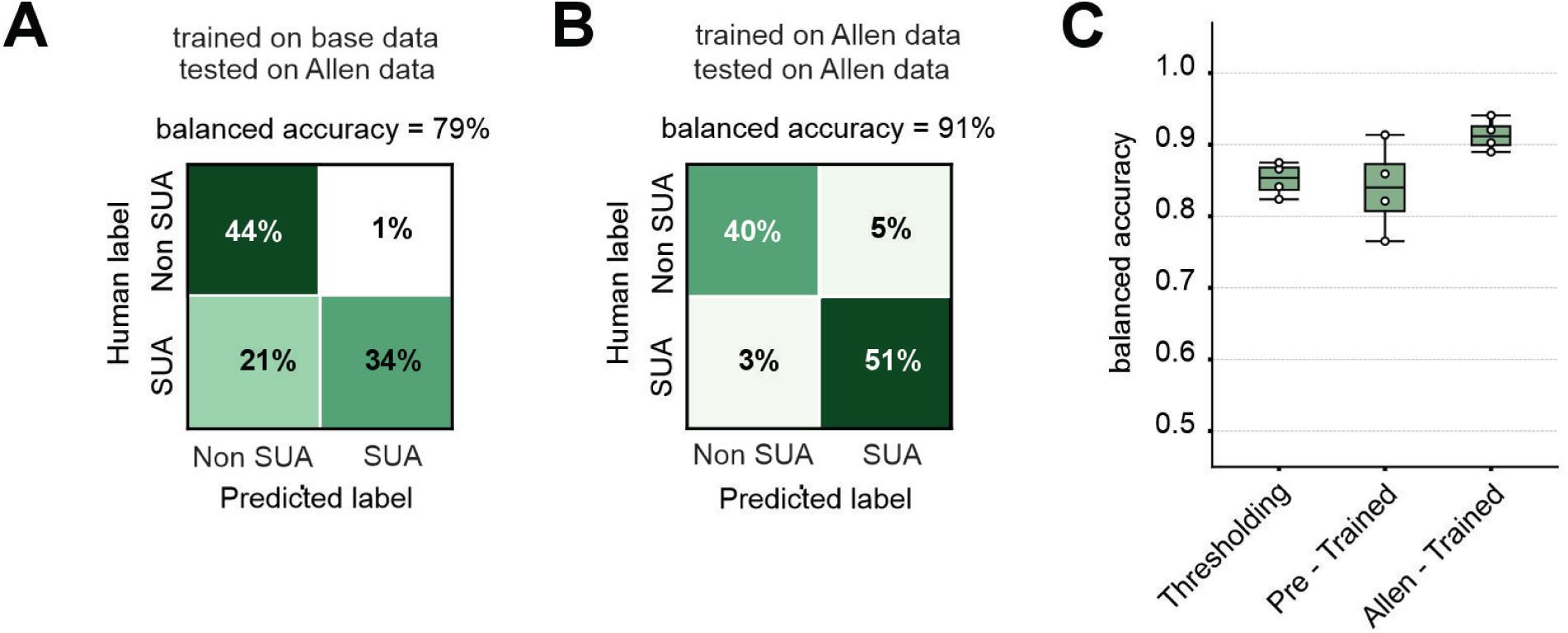
Cross-dataset evaluation of UnitRefine models trained on the base dataset and re-trained on the Allen dataset. (**A**) Confusion matrix showing the performance of the model trained on the base dataset, achieving a balanced accuracy of 79%. (**B**) Confusion matrix showing the performance of the model retrained on the Allen dataset, achieving a higher balanced accuracy of 91%. (**C**) Comparison of balanced accuracy across different approaches: a threshold-based curation method, the pre-trained model (base dataset), and the retrained model (Allen dataset). The retrained model on the Allen dataset outperforms both the pre-trained and threshold-based approaches, demonstrating improved generalization.

**Figure S5:**
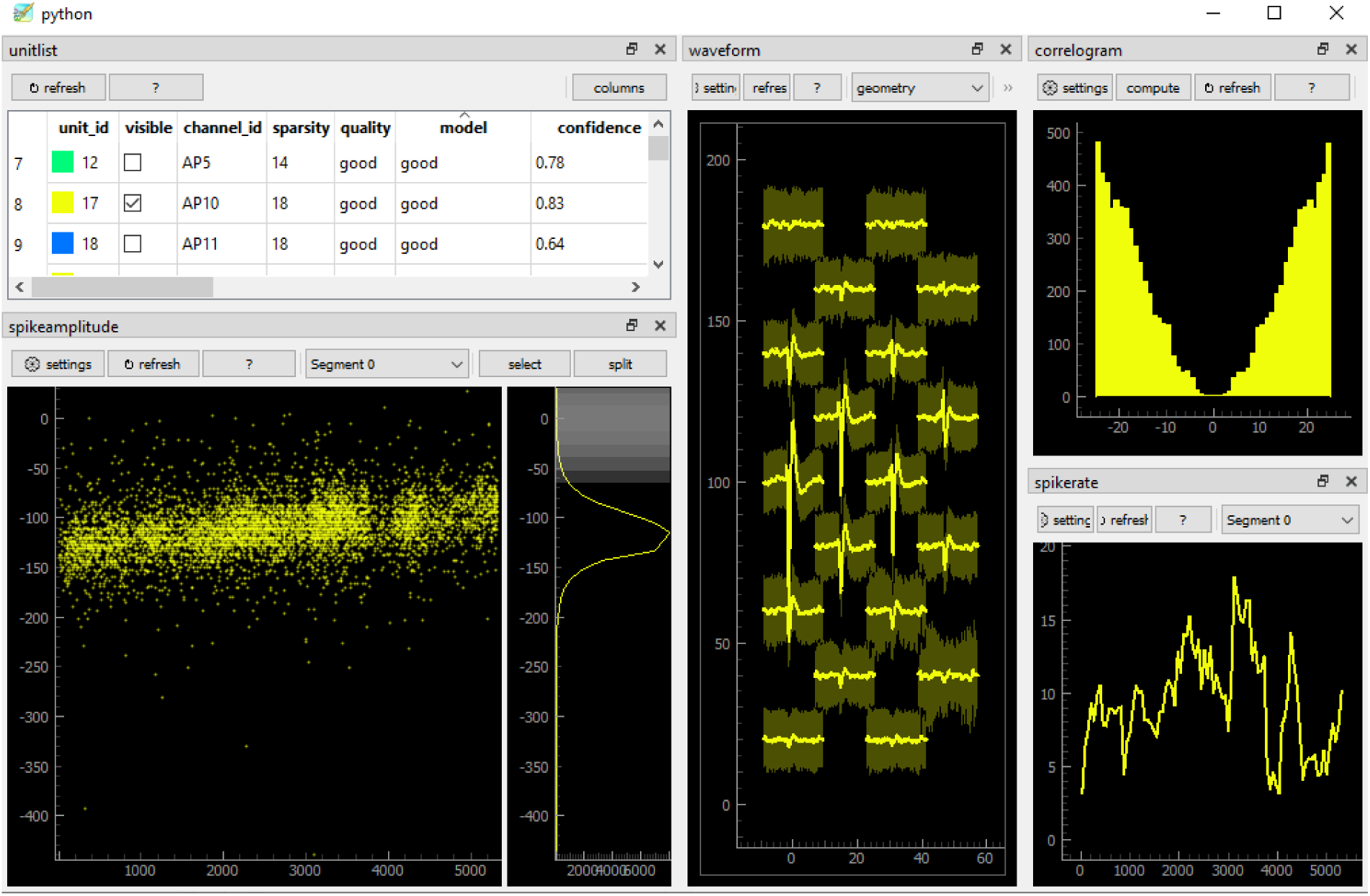
UnitRefine curation interface. Cluster inspection is performed through the SpikeInterface-GUI backend, which displays waveforms, amplitudes, correlograms, and firing rates. Users can manually add or refine labels using hotkeys, with assigned labels shown in the quality column.

**Figure S6:**
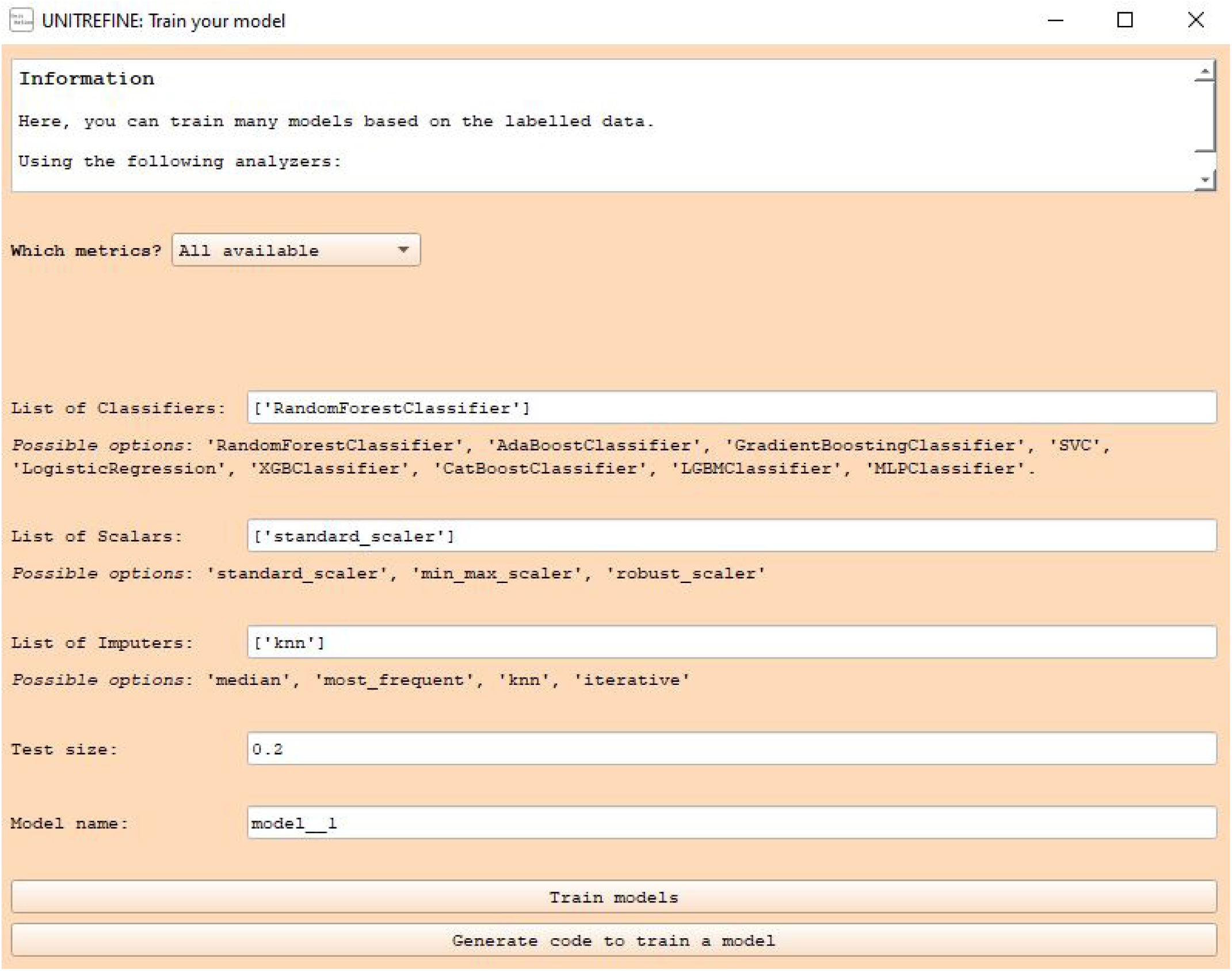
Training a model with the UnitRefine GUI. The interface allows users to select metrics, classifiers, scalers, and imputers, specify training parameters, and train new models from curated labels. It also supports automatic code generation for reproducible model training.

## Footnotes

1 https://github.com/cortex-lab/phy/

2 https://github.com/anoushkajain/UnitRefine/tree/main/UnitRefine/dataset

3 https://spikeinterface.readthedocs.io/en/latest/tutorials/qualitymetrics/plot_3_quality_metrics.html

4 https://spikeinterface.readthedocs.io/en/latest/modules/postprocessing.html

5 https://huggingface.co/AnoushkaJain3/UnitRefine-mice-sua-classifier

6 https://github.com/int-brain-lab/paper-brain-wide-map/blob/main/brainwidemap/meta/region_info.csv

7 https://github.com/anoushkajain/UnitRefine

8 https://spikeinterface.readthedocs.io/en/stable/modules/preprocessing.html

9 https://github.com/billkarsh/CatGT

10 https://github.com/kwikteam/phy

11 https://spikeinterface.readthedocs.io/en/latest/modules/qualitymetrics.html

